# Thalamocortical credit routing from the basal ganglia and cerebellum: anatomical constraints and circuit principles

**DOI:** 10.64898/2026.10.02.755501

**Authors:** Sho Tanno, Mieko Morishima, Kyohei Fujita, Shigeki Kato, Kazuto Kobayashi, Yasuo Kawaguchi, Yoshikazu Isomura, Riichiro Hira

## Abstract

Animals learn to select actions that maximize reward in complex environments. A central challenge in this process is credit routing: behavioral outcomes must be routed back to the neural populations that contributed to generating the relevant actions. One possible route is through thalamocortical pathways conveying learning signals from the basal ganglia and cerebellum, where plasticity is shaped by reward-prediction errors. However, how these pathways route such signals back to specific cortical circuits remains unclear. To address this question, we combined anatomical circuit mapping in mice with closed-loop circuit modeling. Anterograde transsynaptic tracing revealed that thalamic neurons receiving input from the substantia nigra pars reticulata or the lateral cerebellar nucleus were largely segregated, with partial overlap in the ventromedial thalamus. Nevertheless, their cortical axons shared a pronounced enrichment in layer 1a, most prominently in secondary motor cortex. In a brain-wide synergistic loop-circuit model, learning was enhanced when both pathways delivered additive and gain-modulating feedback aligned with the cortical action readout. Local activity-based calibration established this alignment, whereas dendritic-spike- and reward-prediction-error-dependent thalamocortical plasticity further refined behavioral output. Module-specific feedback was essential for a brain–machine interface task requiring the simultaneous conditioning of multiple target neurons. Together, these findings define a credit-routing architecture in which laminar convergence, plastic alignment, and modular feedback link parallel subcortical learning systems to cortical action representations.

## Introduction

Reinforcement learning requires updating action values based on behavioral outcomes while directing circuit reorganization specifically to the neural populations that generated the relevant actions. The basal ganglia and cerebellum are two major subcortical systems implicated in these processes. Both receive cortical input and return processed signals to the cortex through the thalamus, forming recurrent loops that support motor control, action selection, timing, and reward-dependent learning (Alexander et al., 1986; Bostan and Strick, 2018; Schultz et al., 1997; Reynolds et al., 2001; Samejima et al., 2005; Williams and Eskandar, 2006; Kravitz et al., 2012; Tai et al., 2012; Steinberg et al., 2013; Yagishita et al., 2014; Wagner et al., 2017; Kostadinov et al., 2019; Larry et al., 2019; Sendhilnathan et al., 2020; Nonomura et al., 2018; Larry et al., 2024; Jin and Hull, 2025). However, it remains unclear which cortical populations receive these outputs and how the returning signals modify cortical computation. Understanding learning in these systems therefore requires determining the pathways through which learning-related changes in the basal ganglia and cerebellum are returned to the cortex.

The basal ganglia and cerebellum have classically been viewed as functionally segregated systems operating through largely parallel channels. Indeed, in primates, transsynaptic tracing revealed topographically organized pathways linking the two systems with partly distinct cortical areas (Hoover and Strick, 1999; Kelly and Strick, 2003). In rodents, basal ganglia output preferentially targets the ventromedial nucleus (VM) and the rostroventral portion of the ventral anterior–ventral lateral complex (VA–VL), whereas large cerebellar terminals are concentrated in the caudodorsal VA–VL region (Kuramoto et al., 2011). Moreover, VM and the rostroventral region of VA–VL project broadly and preferentially to cortical layer 1 (L1), whereas neurons in the caudodorsal region of VA–VL preferentially innervate the middle layers of more restricted cortical areas (Shigematsu et al., 2016; Ushimaru et al., 2012; Kuramoto et al., 2009, 2015; Tanaka et al., 2018). Thus, at the level of the motor thalamus and the target cortical layers, basal ganglia and cerebellar signals remain largely segregated.

In contrast, recent anatomical and physiological studies have revealed a less sharply segregated organization. Large-scale tracing studies showed that both basal ganglia-related and cerebellar loops extend broadly across sensorimotor and association cortices (Foster et al., 2021; Pisano et al., 2021). Within the mouse ventral motor thalamus, cerebellar axons extend across multiple nuclei, including regions enriched in basal ganglia terminals (Alonso-Martínez et al., 2023). Electrophysiological analyses further demonstrated that inputs from the GPi/SNr and deep cerebellar nuclei can converge onto individual motor-thalamic neurons (Koster and Sherman, 2024). Cerebellar output is itself distributed across multiple thalamic routes. Fastigial nucleus neurons innervate several thalamic nuclei (Fujita et al., 2020), and a lateral cerebellar nucleus (LCN)–thalamus–secondary motor cortex (M2) pathway conveys learning-related, reward-dependent post-error signals to L1 (Ako et al., 2025). How basal ganglia and cerebellar outputs converge in M2 is particularly important, given its correspondence to premotor and supplementary motor areas in primates and its central role in reward-guided learning and action selection (Hira et al., 2013a,b; Sul et al., 2011; Wang et al., 2020b; Augusto et al., 2025; Mushiake et al., 1991; Hoshi & Tanji, 2000). Thus, it remains unclear to what extent basal ganglia and cerebellar signals are segregated or integrated within the motor thalamus and whether this organization is preserved or reorganized across their cortical target layers.

The laminar destination of these signals is particularly important because L1 input can influence cortical pyramidal neurons in a fundamentally different manner from input to the middle and deep layers. L1 contains the distal apical dendrites of pyramidal neurons, where long-range inputs can interact with proximal synaptic drive and backpropagating action potentials to evoke dendritic Ca²⁺ spikes and burst firing (Rubio-Garrido et al., 2009; Larkum et al., 1999, 2004; Xu et al., 2012). Dendritic Ca²⁺ spikes and plateau potentials can, in turn, gate or induce long-lasting synaptic potentiation, providing a direct link between dendritic coincidence events and cortical plasticity (Kampa et al., 2006; Gambino et al., 2014; Cichon and Gan, 2015). Such nonlinear dendritic integration makes L1 input well suited to regulate neuronal gain and the induction of plasticity. By contrast, thalamic inputs terminating in the middle and deep layers are positioned to act more directly on proximal dendritic compartments and local cortical circuits, thereby contributing more directly to postsynaptic drive rather than primarily through distal-dendritic gain modulation (Hooks et al., 2013). Laminar targeting is therefore a critical determinant of how basal ganglia and cerebellar outputs influence cortical computation and learning.

When thalamocortical pathways convey learning-related signals generated in the basal ganglia and cerebellum, not only laminar specificity but also the specificity of the cortical cell populations they influence becomes critically important. Learning-related signals generated in the basal ganglia and cerebellum must return to the cortical populations whose activity contributed to the selected action. We refer to this circuit-level assignment of learning signals through closed loops as credit routing. Without appropriate credit routing, dopamine-dependent synaptic plasticity in the striatum and climbing-fiber-instructed plasticity at Purkinje cell synapses would not necessarily lead to appropriate reorganization of the cortical circuits responsible for generating the relevant actions. Plasticity of thalamocortical pathways during motor learning may therefore play a key role in establishing appropriate credit routing (Biane et al., 2016; Tanaka et al., 2018; Ramot et al., 2025). At the same time, three-factor learning rules in the cortex that combine presynaptic activity, a postsynaptic dendritic event, and a reward-related modulatory signal could use reward-related information available to the basal ganglia and cerebellum to coordinate plasticity across these three systems (Frémaux and Gerstner, 2016; Gerstner et al., 2018).

Our previous synergistic reinforcement-learning model showed how the basal ganglia and cerebellum, despite their distinct response dynamics and reward-prediction-error-dependent plasticity, can cooperate over time (Yoshida et al., 2025). That model focused on the temporal aspect of cooperation between the two systems but did not address how their outputs are routed through the thalamus back to the cortical populations responsible for action. The demands placed on circuits for credit routing are substantial because behaviorally relevant information is encoded across cortical populations in a spatially and temporally distributed manner (Nelson et al., 2021). This problem becomes particularly evident not only in conventional operant learning but also in neural operant conditioning and brain–machine interface learning, in which selected cortical populations can be independently reinforced (Fetz, 1969; Hira et al., 2014; Clancy et al., 2014; Francioni et al., 2026).

To address how subcortical learning systems are linked to specific cortical populations through the thalamus, we investigated two questions. First, using anatomical tracing, we examined how basal ganglia and cerebellar outputs are segregated or integrated within the thalamus and across cortical layers, with particular emphasis on thalamic pathways targeting M2. Second, we integrated these anatomical and physiological findings into a computational model to determine how the organization and plasticity of these thalamocortical pathways can implement credit routing. Thus, our study identifies the anatomical requirements and computational sufficiency conditions that enable biologically implemented credit routing and provides a new framework in which the cerebral cortex, basal ganglia, and cerebellum are integrated through the thalamus to support coordinated reinforcement learning across the brain.

## Results

### Retrograde tracing identifies basal ganglia and cerebellar inputs to thalamic nuclei projecting to M2

To identify the subcortical pathways that reach M2 through the thalamus, we first mapped the thalamic nuclei connected with M2 by co-injecting the anterograde tracer biotinylated dextran amine (BDA) and the retrograde tracer cholera toxin subunit B (CTB) into M2 (**Fig. 1a,b**). Immunostaining for neurofilament 200 (N200) delineated the M1–M2 boundary and verified the location of the injection site (**Fig. 1c**). In the thalamus, BDA-labeled corticothalamic axons and CTB-labeled thalamocortical neurons were observed in overlapping regions, indicating reciprocal connectivity between M2 and the thalamus (**Fig. 1d–g**). GAD67 and calbindin immunostaining localized CTB-positive neurons primarily to the ventral anterior (VA), ventrolateral (VL), ventromedial (VM), and posterior (Po) nuclei, consistent with previous anatomical findings (Ushimaru et al., 2012). Abbreviations of thalamic nuclei and neighboring regions are listed in **Extended Data Table 1.**

**Figure 1.**
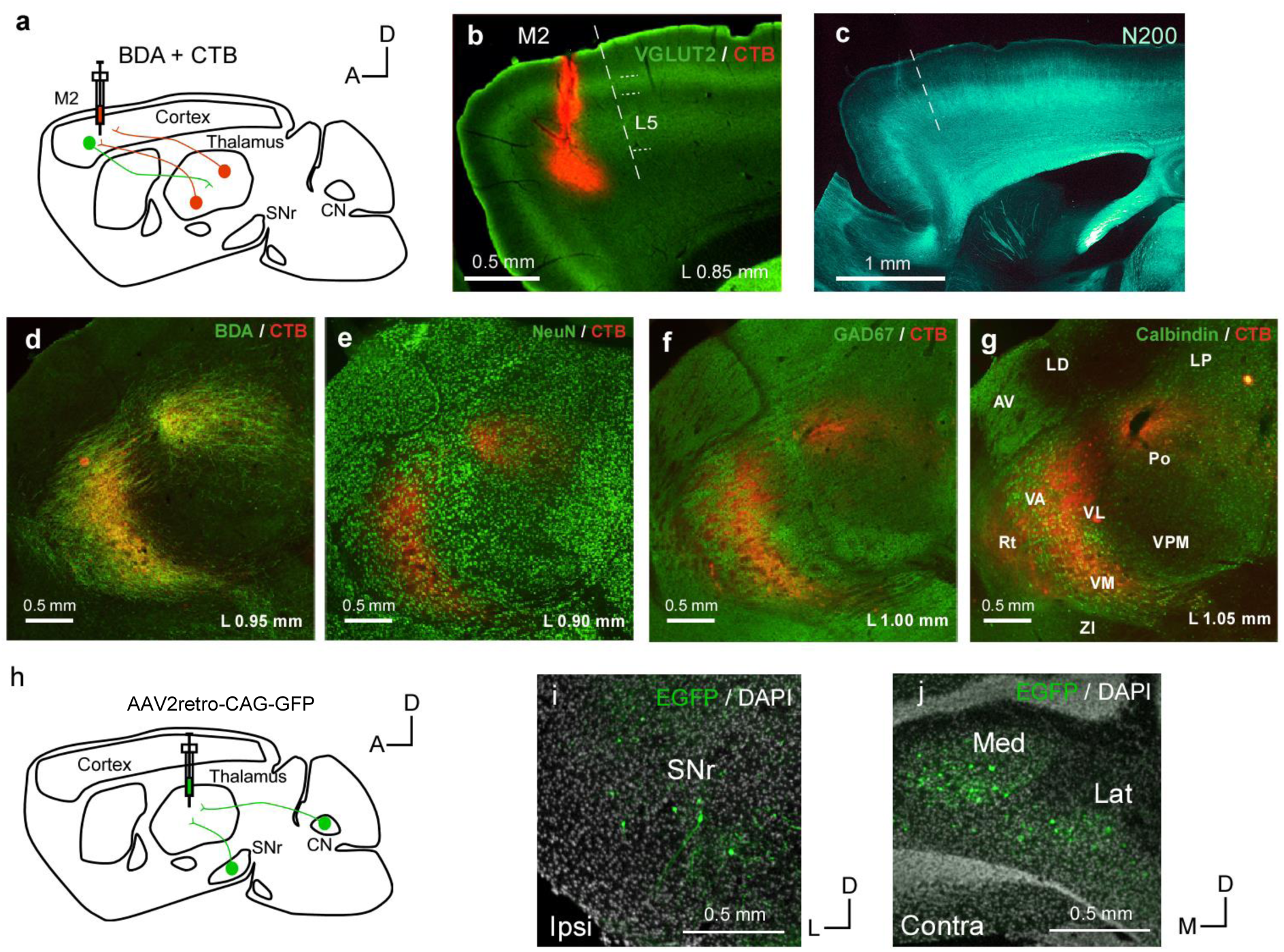
| Reciprocal connectivity between M2 and motor thalamus and subcortical inputs to M2-projecting thalamic regions **a,** Schematic of the injection of BDA and the retrograde tracer CTB-Alexa 555 into M2. **b,** CTB injection site in M2 (red) and VGLUT2 immunostaining (green). Dashed lines indicate cortical layer boundaries. Sagittal section 0.85 mm lateral to the midline. Scale bar, 0.5 mm. **c,** Sagittal section stained for Neurofilament 200 (N200). Dashed lines indicate the boundary between M1 and M2. Scale bar, 1 mm. **d,** Overlay of CTB labeling (red) and BDA labeling (green) in the thalamus. **e,** Overlay of CTB labeling (red) and NeuN immunostaining (green) in the thalamus. **f,** Overlay of CTB labeling (red) and GAD67 immunostaining (green) in the thalamus. **g,** Overlay of CTB labeling (red) and Calbindin immunostaining (green) in the thalamus. Thalamic nuclear boundaries were identified on the basis of these markers and anatomical landmarks. **h,** Schematic of the injection of AAV2retro-CAG-GFP into thalamus to retrogradely label subcortical neurons projecting to the injected thalamic nuclei. **i,** Representative coronal image of retrogradely labeled cells in the ipsilateral SNr. **j,** Representative coronal image of retrogradely labeled cells in the contralateral CN, including the medial (Med) and lateral (Lat) nuclei.

We next injected AAV2retro-CAG-GFP into these M2-projecting thalamic nuclei to identify the subcortical neurons projecting to them (**Fig. 1h–j**). Retrogradely labeled neurons were found in the ipsilateral substantia nigra pars reticulata (SNr) and the contralateral cerebellar nuclei, including the medial and lateral cerebellar nuclei. These distributions were consistent with previous descriptions of basal ganglia and cerebellar projections to the motor thalamus (Kuramoto et al., 2011; Gornati et al., 2018; Alonso-Martínez et al., 2023). Thus, thalamic nuclei projecting to M2 receive inputs from both basal ganglia and cerebellar output structures, establishing the anatomical pathway examined by input-defined anterograde transsynaptic tracing in the subsequent experiments.

### SNr and LCN inputs target largely distinct thalamic populations with restricted overlap in VM

We next used high-titer AAV1-mediated anterograde transsynaptic labeling to identify thalamic neurons receiving input from the SNr or LCN and to visualize their cortical axons (**Fig. 2a–f**; Zingg et al., 2017, 2020). AAV1-FlpO was injected into either the ipsilateral SNr or the contralateral LCN, and the FlpO-dependent reporter AAV-CAG-FRT-Rev-tdTomato was injected into the thalamus. This strategy induced tdTomato expression in thalamic neurons receiving input from the injected subcortical nucleus, thereby labeling both their somata in the thalamus and their axonal projections in the cerebral cortex. Because the present study focused on cerebellar output to higher-order motor cortex, we analyzed the LCN as the principal cerebellar source. SNr-recipient thalamic neurons were located mainly in and around VM, with additional labeling near Po and CL, whereas LCN-recipient neurons were concentrated primarily in VL (**Fig. 2b–f**). Exploratory studentized residual-permutation tests detected source-by-nucleus interactions for normalized cell count (p=0.040), cell density (p=0.0055), and normalized cell density (p=0.029; permutation tests; **Fig. 2g,h).** Thus, the two input-defined neuronal populations were distributed largely in distinct thalamic nuclei, consistent with previous anatomical studies (Kuramoto et al., 2011; Gornati et al., 2018; Alonso-Martínez et al., 2023).

**Figure 2.**
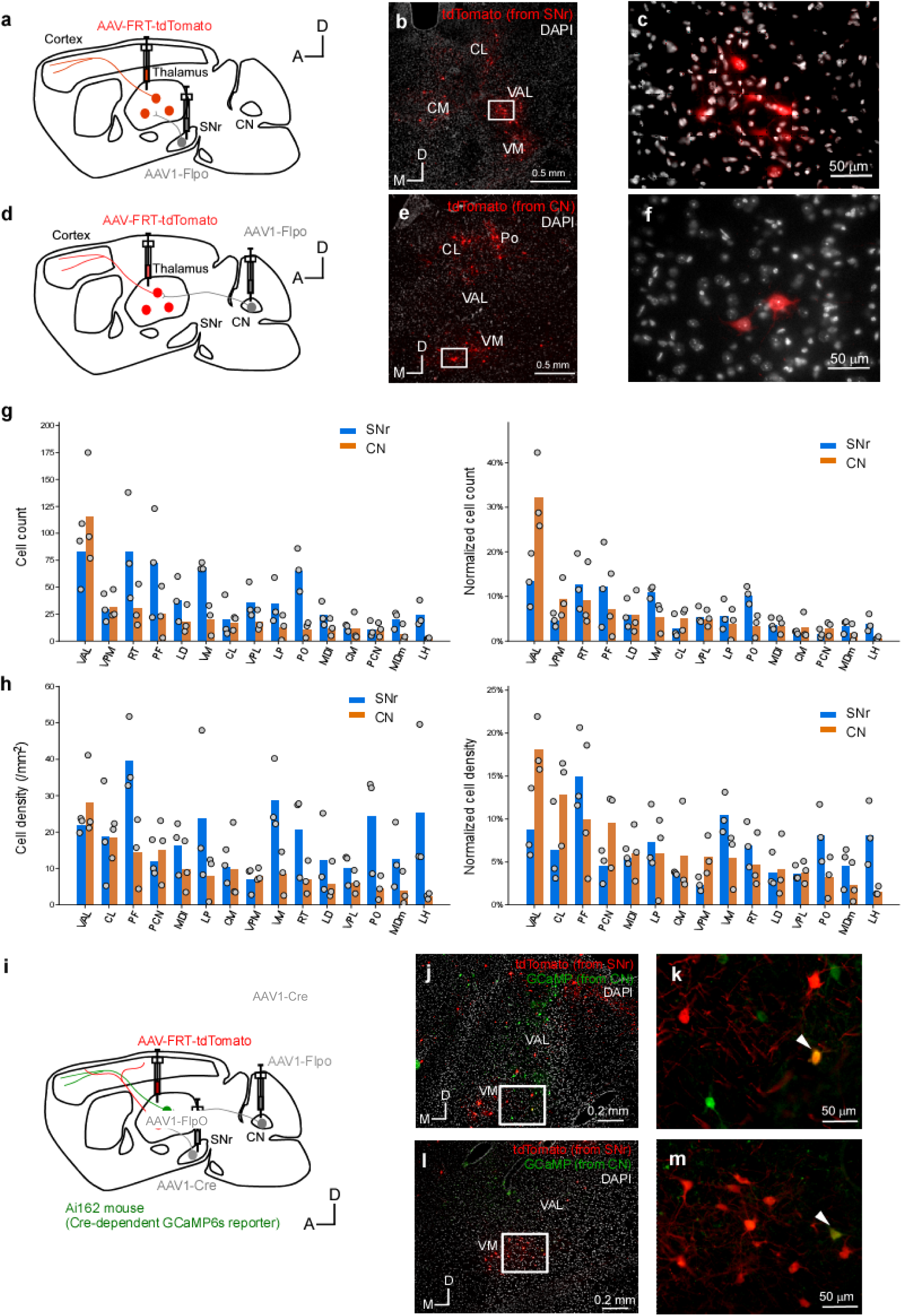
| SNr- and LCN-recipient thalamic populations are largely segregated with partial overlap in VM. a, Strategy for labeling SNr-recipient thalamic neurons using AAV1-FlpO in the SNr and a FlpO- dependent tdTomato reporter in the thalamus. **b,** Representative coronal section showing tdTomato-labeled thalamic neurons (red) and DAPI (white). **c,** Enlargement of the boxed region in **b.d,** Corresponding strategy for labeling LCN-recipient thalamic neurons. **e,** Representative coronal section showing labeled thalamic neurons. **f,** Enlargement of the boxed region in e. **g,** Cell counts (left) and normalized cell count (right) across thalamic nuclei. The SNr and LCN groups (labeled CN in the plots) differed in normalized cell count but not in cell count (studentized Freedman– Lane permutation test: p=0.040 and 0.099, respectively). Quantification in g,h included 2,108 labeled thalamic cells across 42 unique section records from 3 mice in the SNr group and 1,174 cells across 39 unique section records from 3 mice in the LCN group. **h,** Cell densities (left) and normalized cell densities (right) across thalamic nuclei. The SNr and LCN groups differed in both measures (studentized Freedman–Lane permutation test: p=0.0055 and 0.029, respectively). **i,** Dual-labeling strategy in an Ai162 reporter mouse using orthogonal Cre- and FlpO-dependent labeling to distinguish SNr-recipient and LCN-recipient thalamic populations. **j,** Distribution of SNr-defined labeling (red) and LCN-defined labeling (green) in motor thalamus. **k,** Enlargement of the boxed region in **j.** Arrowheads indicate double-labeled neurons. **l,m,** Corresponding images for **j** and **k** from a second mouse.

To test whether the two populations nevertheless overlapped locally, we labeled both pathways in the same animal using two recombinase-dependent labeling strategies **(Fig. 2i– m).** To minimize potential competition between two reporter AAVs in the thalamus, which could lead to underestimation of cellular overlap (de Alencastro et al., 2020), we used a fluorescent reporter mouse and injected only one reporter AAV into the thalamus. Green fluorescence in thalamic neurons was produced by GCaMP6s expression in Ai162 mice (Cre-dependent GCaMP6s reporter mice; Daigle et al., 2018), whereas red fluorescence was produced by AAV-CAG-FRT-Rev-tdTomato. Neurons coexpressing GCaMP6s and tdTomato were observed in VM, demonstrating local cellular overlap between SNr- and LCN-recipient populations despite their broader segregation across the motor thalamus.

Together, these results define a two-level organization: broad separation of basal ganglia- and cerebellar-recipient populations across the motor thalamus, accompanied by restricted cellular overlap within VM.

### SNr- and LCN-recipient thalamic neurons share an L1a-enriched cortical projection component, most prominently in M2

Although basal ganglia and cerebellar outputs projected predominantly to distinct thalamic nuclei, it remained unclear whether these pathways would project to different cortical layers or converge within the same layers. To address this question, we first examined the cortical layers innervated by thalamic neurons receiving SNr input (**Fig. 3a–c**). In M2, their axons exhibited a clear distribution in L1a, together with projections to the deeper part of L2/3. In M1, they showed a strong fluorescence peak in L1, particularly L1a, as well as fluorescence in deep L2/3. In S1, fluorescence was observed in L1a and deep L6. Thus, L1 was one of the major cortical projection targets of thalamic neurons receiving SNr input across multiple cortical areas.

**Figure 3.**
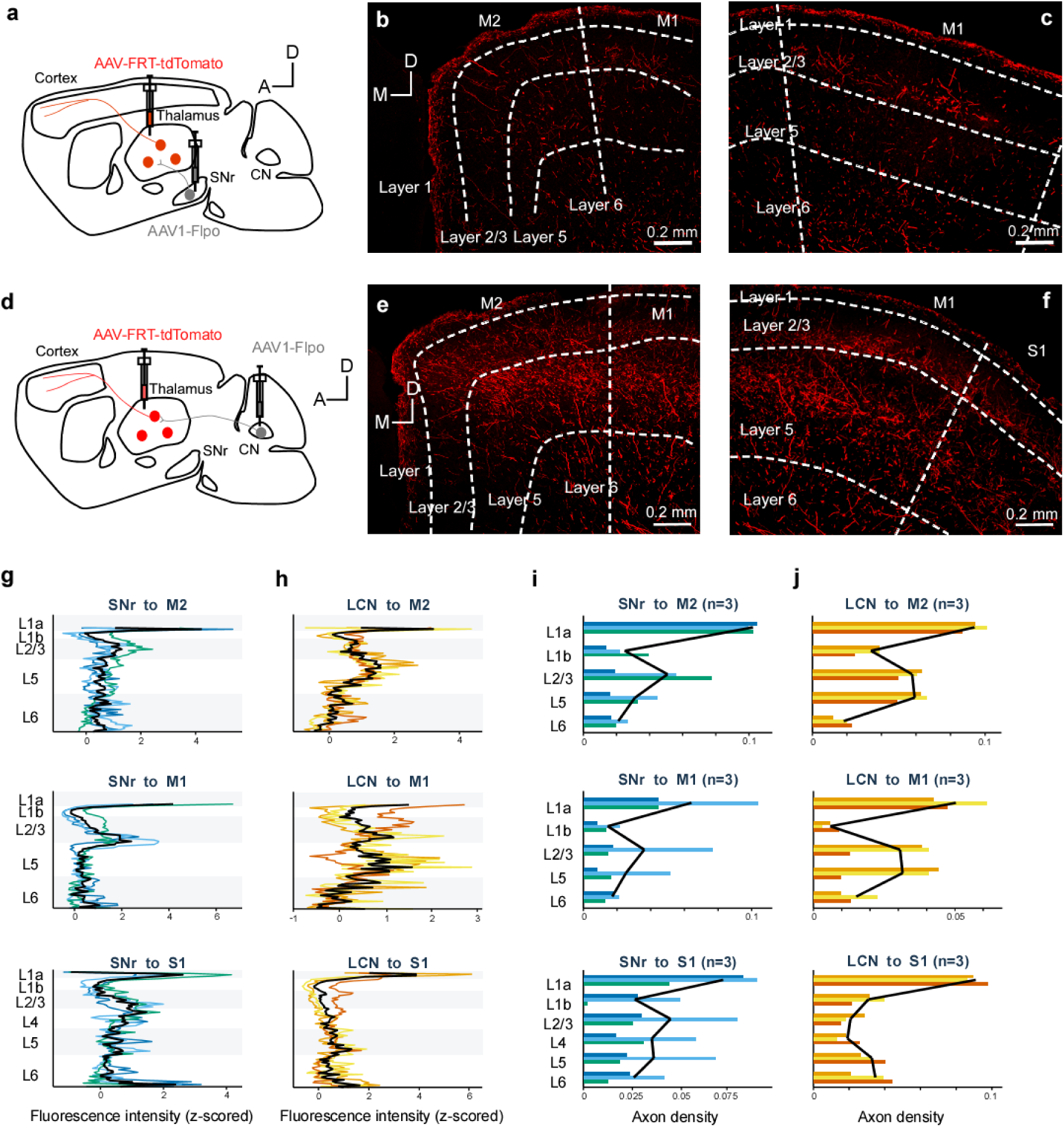
| Laminar organization of cortical projections from SNr- and LCN-recipient thalamic neurons. a, Strategy for labeling SNr-recipient thalamic neurons and their cortical axons. **b,** Representative labeled axons (red) spanning M2 and M1. Dashed lines indicate cortical areas and layers. **c,** Representative labeled axons in M1. **d,** Corresponding strategy for labeling LCN-recipient thalamic neurons. **e,** Representative labeled axons spanning M2 and M1. **f,** Representative labeled axons spanning M1 and S1. **g,** Background-subtracted, z-scored fluorescence profiles across cortical depth for the SNr-recipient pathway. Colored lines represent individual mice; black lines represent the mean. **h,** Corresponding profiles for the LCN-recipient pathway. **i,** Layer-wise axon-length measure for the SNr-recipient pathway. Bars represent individual mice and black lines the mean (n = 3 mice). **j,** Corresponding layer-wise axon-length measure for the LCN-recipient pathway (n = 3 mice).

We next examined the cortical layers innervated by thalamic neurons receiving input from the LCN (Fig. 3d–f). Because LCN-recipient cells were located mainly in VL, the classical laminar- segregation model would predict preferential projections to the middle cortical layers. However, analysis of fluorescence intensity across cortical depth in M1, M2, and S1 revealed clear L1 signals from axons of LCN-recipient thalamic neurons in all three areas. In M2, axons were distributed in L1a, L2/3, and L5. In M1, fluorescence was detected in L1a as well as L2/3 and L5. In S1, a strong distribution was observed in L1a. Thus, the thalamocortical pathway arising from LCN-recipient cells strongly innervated both L1 and the middle or deep layers in M1 and M2, and predominantly L1 in S1.

In contrast, the pathway from the medial cerebellar nucleus (MCN) through the thalamus to the cerebral cortex showed little projection to M1, and the sparse M1 projection did not exhibit L1 predominance (**Extended Data Fig. 1a–c**).

To compare these results across animals, we quantified both fluorescence intensity across cortical depth and total axon length within each layer in three mice per group (**Fig. 3g–j**). In M2, M1, and S1, both the SNr and LCN pathways showed their largest projection component in L1a. These findings suggest that basal ganglia and cerebellar outputs routed through the SNr and LCN, respectively, exhibit similar cortical projection patterns, particularly in L1.

### Integrated public datasets recapitulate an upper-L1-enriched return pattern

We next used publicly available anatomical resources to perform virtual transsynaptic tracing (**Fig. 4a**). Bulk anterograde AAV tracing data for the SNr and cerebellar nuclei (CN) were obtained from the Allen Mouse Brain Connectivity Atlas (Oh et al., 2014). The CN group included experiments in which the lateral cerebellar nucleus, annotated as the dentate nucleus in the Allen atlas, was a primary or secondary injection site. Experiments selectively targeting inhibitory CN neurons were excluded. Whole-brain single-neuron axonal reconstructions acquired using fluorescence micro-optical sectioning tomography (fMOST) were obtained from the Brain Image Library (BIL) collection generated as part of the BRAIN Initiative Cell Census Network (BICCN) (Peng et al., 2021). The Allen dataset provided the distribution of SNr- and CN-derived projection density across individual thalamic nuclei (**Fig. 4b**), whereas the BIL/BICCN dataset provided the axon length in each cortical layer for neurons assigned to individual thalamic nuclei (**Fig. 4c,d**). We assumed that axons projecting to the thalamus innervated individual thalamic neurons non-selectively. Under this assumption, the weighted average of the projection density from the SNr or CN to each thalamic nucleus and the layer-specific cortical projection density of neurons within that nucleus should approximate the cortical laminar projection density associated with the SNr or CN. We termed this virtual transsynaptic tracing approach **Bulk-Single Virtual Tracing (BSVT)**.

**Figure 4.**
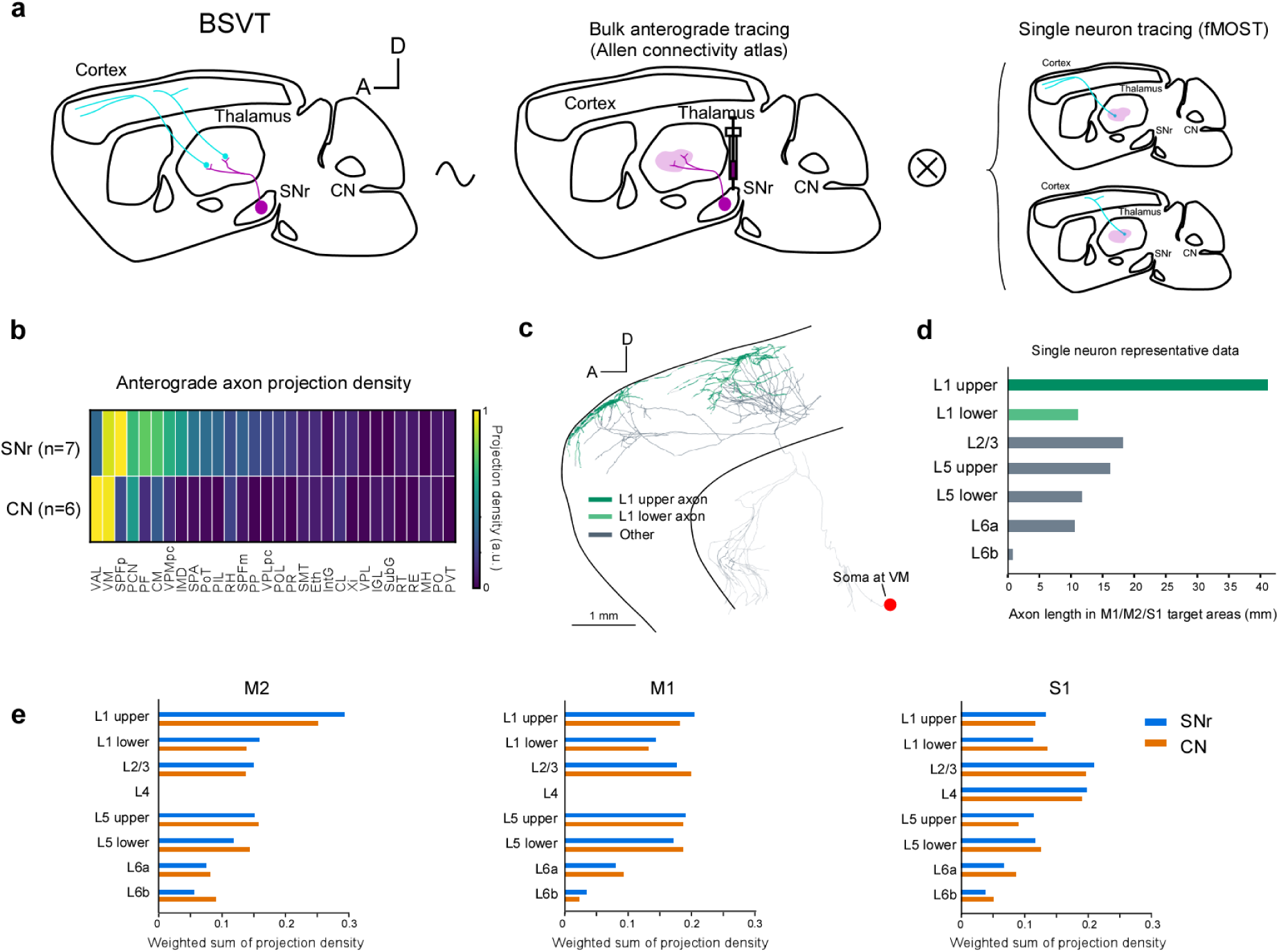
| Integrated anatomical analysis estimating the laminar distribution of thalamocortical projections associated with the basal ganglia and cerebellum. a, Overview of Bulk-Single Virtual Tracing (BSVT). Virtual transsynaptic tracing was performed by integrating anterograde tracing data from the Allen Mouse Brain Connectivity Atlas with fMOST single- neuron tracing data. **b,** Axonal projections from the SNr (n = 7) or CN (n = 6) within individual thalamic nuclei. Projection density (the fraction of detected projection pixels) was divided by injection volume; each source column was then scaled by its maximum for display. In an exploratory analysis of the five nuclei with the highest pooled normalized projection density, the relative projection pattern differed (PERMANOVA, p=5.83×10^-4^). **c,** Reconstruction of a representative single thalamic neuron with its soma in VM. Axons are classified as projecting to upper L1, lower L1, or other layers. **d,** Layer-specific axon length within cortical targets for the representative neuron shown in **c**. **e,** BSVT results showing layer fractions of weighted projection density of M2, M1, and S1. The distributions differed significantly in S1, but not in M2 or M1 (Holm-adjusted exact PERMANOVA: M2, p=0.090; M1, p=0.090; S1, p=0.007).

BSVT revealed upper-L1 predominance in M2 for both the SNr and CN pathways (**Fig. 4e**). The two pathways also showed similar laminar specificity in M2, M1, and S1. However, the SNr pathway showed a systematic tendency toward more superficial projections, whereas the cerebellar pathway tended to project more deeply. These results were largely consistent with our experimental anterograde transsynaptic tracing.

BSVT used single-neuron axonal tracing data to characterize thalamocortical projections. Because the Allen Mouse Brain Connectivity Atlas also provides bulk AAV injection data for thalamocortical projections, we performed a corresponding analysis using bulk data for both the subcortical-to-thalamic and thalamic-to-cortical projections. We termed this method **Bulk- Bulk Virtual Tracing** (**BBVT; Extended Data Fig. 2a**). In the BBVT analysis, both the SNr and cerebellar pathways showed L6 predominance (**Extended Data Fig. 2b**).

To further examine the difference between BSVT and BBVT, we compared the density of thalamocortical axons with the average laminar specificity of thalamocortical projections estimated from the databases. The former was assessed using the fluorescence density of VGLUT2 immunostaining, which is concentrated at thalamocortical axon terminals (**Extended Data Fig. 2c**). For the latter, output densities from the 12 matched thalamic nuclei were averaged with equal weight for both BSVT and BBVT (Extended Data Fig. 2d). BSVT closely matched the distribution of thalamocortical axons, whereas BBVT underestimated superficial projections and was biased toward deep layers. This descriptive agreement in the sampled M2 profile supports the superficial component estimated by BSVT, but does not establish population-level equivalence.

### An anatomically constrained BWSLC model links parallel subcortical pathways to cortical output

We next examined the computational significance of the observed laminar organization of thalamocortical projections. To represent two candidate effects of L1-enriched and non-L1 return pathways on cortical activity, we modeled the former as gain-modulating feedback and the latter as additive feedback (**Fig. 5a,b**). Outputs from both the basal ganglia and cerebellar pathways returned to the cortex through the thalamus in these two forms.

**Figure 5.**
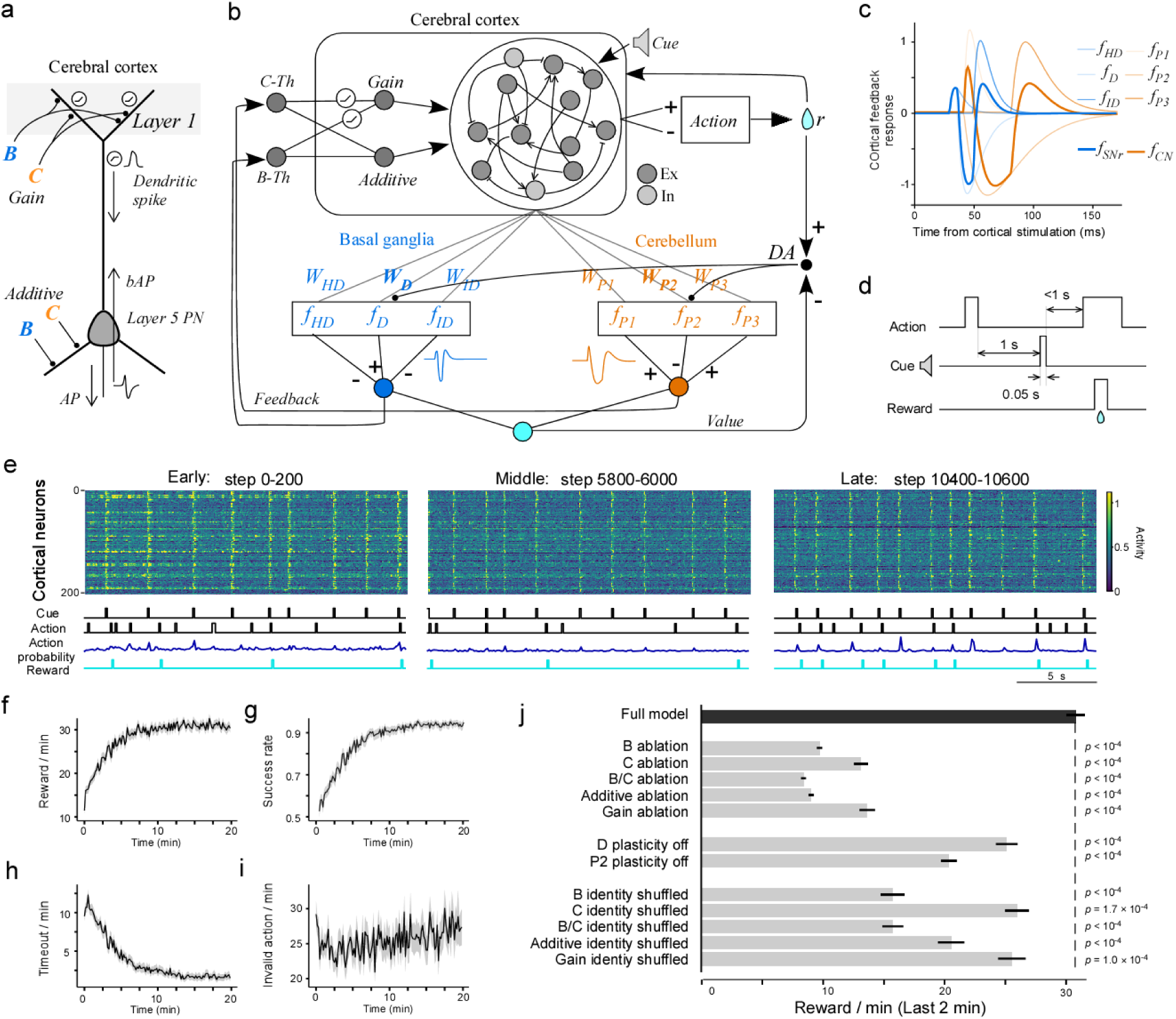
| Cue-triggered action learning in the anatomically constrained BWSLC model. a, Conceptual diagram of a layer 5 projection neuron and layer 1 input. Basal-ganglia-derived signal B and cerebellum-derived signal C induce dendritic activity and dendritic spikes through additive input or gain modulation. bAP, backpropagating action potential; PN, pyramidal neuron; B, basal ganglia; C, cerebellum. **b,** Overall model architecture comprising the cerebral cortex, thalamus, basal ganglia, cerebellum, action selection, value estimation, and dopamine-cell activity. The feedback functions and weights of the basal ganglia and cerebellar pathways are indicated by the symbols shown in the diagram (W). Basal ganglia and cerebellar outputs were divided symmetrically into additive and gain-modulating components that entered the cortex. **c,** Signals evoked by cortical stimulation and returned to the cortex through the thalamus after passing through the basal ganglia pathway (blue) or cerebellar pathway (orange). A common thalamocortical return delay of 20 ms was applied in addition to the pathway-specific delays. **d,** Temporal structure of the simulated task. A Cue was presented 1 s after the end of an Action, and a reward was delivered when an Action occurred 100–1000 ms after Cue onset. **e,** Examples of cortical neuronal activity, Cue presentation, selected Actions, action probability, and reward during early learning (steps 0–200), middle learning (steps 5800–6000), and late learning (steps 10400–10600). **f,** Number of rewards per minute over the course of learning. **g,** Success rate over the course of learning. Mean ± SEM across 100 iterations. **h,** Number of timeouts per minute over the course of learning. Mean ± SEM across 100 iterations. **i,** Number of invalid Actions per minute over the course of learning. Mean ± SEM across 100 iterations. **j,** Number of rewards during the final 2 min in the BC model and under signal-ablation, plasticity-off, and input-identity-shuffle conditions. Mean ± SEM across 100 iterations. P values were obtained by comparison with the Full model using the Wilcoxon signed-rank test.

We incorporated this circuit into the synergistic reinforcement-learning model proposed in our previous study (Yoshida et al., 2025), thereby constructing the **Brain-Wide Synergistic Loop Circuit (BWSLC) model.** From the previous model, we retained temporal-difference reward-prediction error (TD-RPE)-dependent plasticity in the basal ganglia direct pathway and the cerebellar parallel fiber–Purkinje cell pathway, together with the pathway-specific temporal responses. We additionally introduced explicit thalamocortical feedback and a sparse cortical recurrent neural network (RNN) obeying Dale’s principle.

The cortical RNN consisted of 200 units divided into eight modules of 25 units each (**Extended Data Fig. 3**). Each module contained 10 Context neurons, 10 motor-related neurons, and five inhibitory neurons. The motor-related neurons comprised five Action neurons and five No-Action neurons. Context neurons received input from the Cue, reward, Action neurons, and No-Action neurons and sent signals to the basal ganglia and cerebellar pathways. The eight modules jointly contributed to action selection, value, reward, and RPE, whereas feedback from the basal ganglia, cerebellum, and thalamus was provided in parallel to individual modules.

Signals sent from each module to the basal ganglia and cerebellar pathways were returned to the same module through the corresponding thalamocortical feedback channel. We refer to this module-to-module correspondence as **module-specific routing**. Within each module, the thalamocortical feedback weight onto each Action neuron was set in proportion to that neuron’s action-readout weight. We refer to this cell-level correspondence as **readout alignment**. Thus, the BWSLC distinguishes module-specific routing, which determines the destination module of the feedback, from readout alignment, which distributes feedback among Action neurons within that module.

### Both subcortical pathways, their plasticity, and aligned feedback support cue-triggered action learning

We trained the model on a cue-triggered action task (Go/No-Go task). In this task, a Cue was presented after the end of an Action, and an Action performed within 1 s of Cue presentation was rewarded (**Fig. 5d**). Training of the BWSLC increased the number of successful actions per unit time (**Fig. 5e,f**). This improvement reflected both an increase in the success rate per Cue and an increase in the number of Cues presented per unit time, or equivalently a reduction in the number of timeouts. In contrast, the number of unrewarded Actions did not decrease (**Fig. 5g–i)**.

We designated this condition as the Full model (BC in the figures) and examined the contribution of individual circuit components (**Fig. 5j**). Removing the basal ganglia pathway, the cerebellar pathway, or both pathways reduced task performance relative to the Full model. Performance was also reduced when plasticity in either the basal ganglia or cerebellar pathway was disabled. Similarly, removal of either additive feedback or gain-modulating feedback impaired learning.

We next shuffled the destination identity of each feedback component, thereby disrupting the correspondence between its source and target. Shuffling the identity of the basal ganglia pathway, cerebellar pathway, both pathways, additive feedback, or gain-modulating feedback reduced performance relative to the Full model. These results indicate that efficient learning required both subcortical pathways, plasticity within each pathway, both feedback modes, and the routing specificity of the feedback.

### Activity-based local calibration establishes readout alignment, and thalamocortical plasticity further refines behavioral output

In the Full model, the correspondence between each Action neuron’s behavioral readout weight and its thalamocortical feedback weight—that is, readout alignment—was imposed in advance. This correspondence is termed fine tuning (**Fig. 6a**). Learning performance was reduced in the fine-tuning-free condition, in which this correspondence was removed. We therefore asked whether the contribution of each neuron to behavioral output could be approximated from locally available activity information.

**Figure 6.**
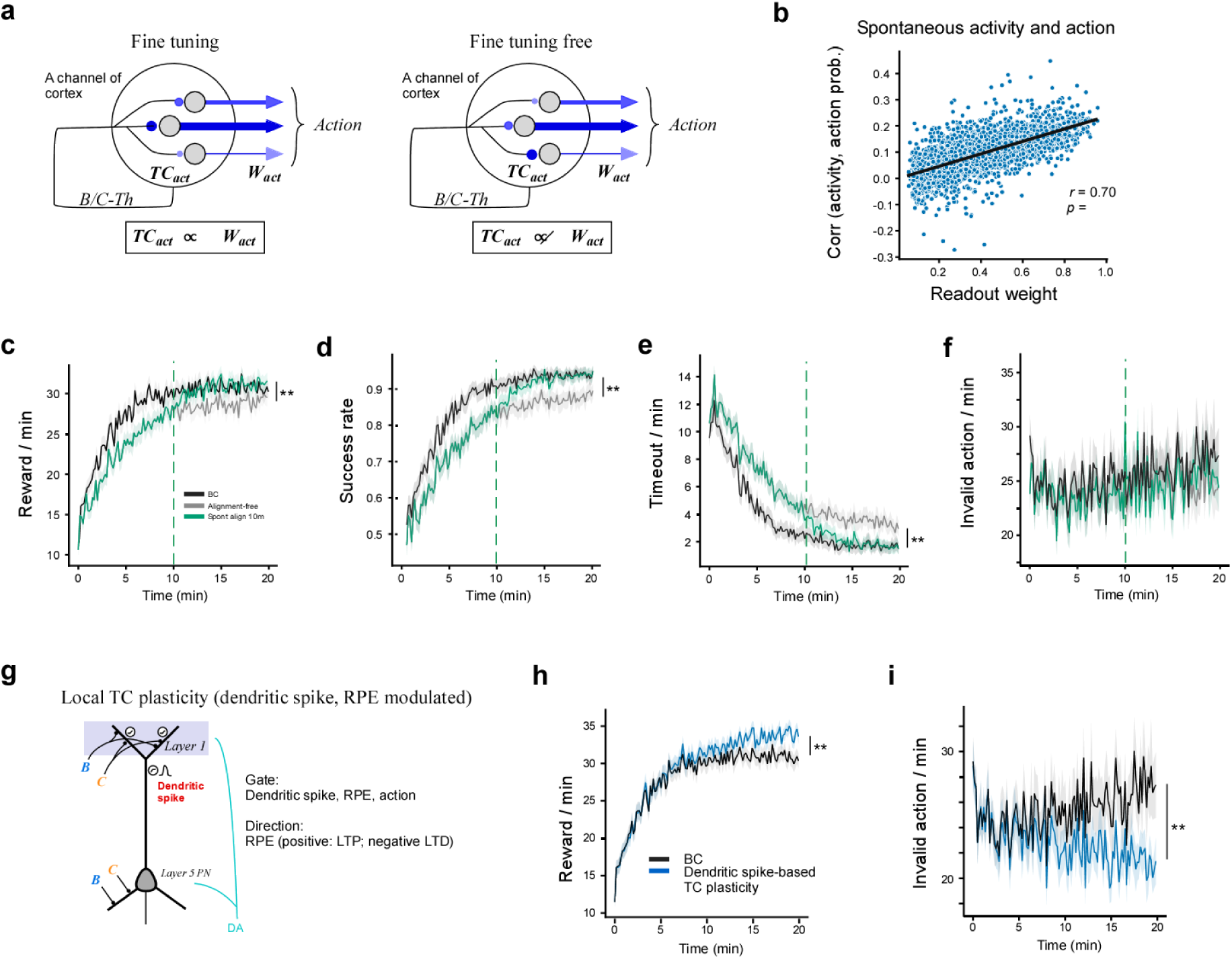
| Activity–action calibration and thalamocortical plasticity restore and refine learning a, Schematic of the fine-tuning condition, in which the behavioral readout weights from cortical activity and the thalamocortical feedback weights were matched, and the fine-tuning-free condition, in which they were not matched. **b,** Correlation between the activity of each Action neuron and action probability during 10 min of spontaneous, task-free behavior, plotted as a function of the neuron’s readout weight (r = 0.70, *p=2.2×10^-593^). Each point represents an individual Action neuron. **c,** Comparison of the BC model, the fine-tuning-free condition, and the condition in which feedback calibrated in a separate 10-min spontaneous run was applied at 10 min of training (local calibration). The reduction in reward number in the fine-tuning-free condition recovered after application of the calibrated feedback. The final 2-min reward rate was higher after local calibration than in the fine- tuning-free condition (Wilcoxon signed-rank test, **p=2.2×10^-3^, n=100). **d,** Time course of the success rate under the conditions shown in **c**. The final 2-min success rate was higher after local calibration than in the fine-tuning-free condition (Wilcoxon signed-rank test, **p=4.0×10^-7^, n=100). **e,** Time course of the number of timeouts per minute under the conditions shown in **c**. The final 2-min timeout rate was lower after local calibration than in the fine-tuning-free condition (Wilcoxon signed- rank test, **p=4.8×10^-7^, n=100). **f,** Time course of the number of invalid Actions per minute under the conditions shown in **c**. The final 2-min invalid-action rate did not differ between the local-calibration and fine-tuning-free conditions (Wilcoxon signed-rank test, p=0.75, n=100). **g,** Schematic of a local thalamocortical plasticity rule gated by dendritic spikes, reward-prediction error (RPE), and the selected action. Positive RPE induced long-term potentiation, whereas negative RPE induced long-term depression. **h,** Time course of the number of rewards in the BC model and in the BC model incorporating dendritic-spike-dependent thalamocortical plasticity (BC+TCP model). The final 2-min reward rate was higher in the BC+TCP model than in the BC model (Wilcoxon signed-rank test, **p=3.3×10^-7^, n=100). **i,** Time course of the number of invalid Actions per minute in the two models. The final 2-min invalid- Action rate was lower in the BC+TCP model than in the BC model (Wilcoxon signed-rank test, **p= 1.6×10^-4^, n=100).

During a 10-min period of spontaneous behavior without task performance, we calculated the correlation between the activity of each Action neuron and the probability of action generation in the model. This correlation coefficient was strongly correlated with the neuron’s actual action-readout weight (r = 0.70; **Fig. 6b**). We therefore trained a model without readout alignment for 10 min, inserted a 10-min spontaneous-behavior period, and updated the thalamocortical feedback weights using the activity–action correlations measured during this period. We refer to this procedure as local calibration.

Following local calibration, the number of rewards per minute recovered within several minutes to a level comparable to that of the Full model with imposed readout alignment (**Fig. 6c**). The success rate and number of timeouts similarly recovered, whereas the number of invalid Actions was comparable to that of the Full model (**Fig. 6d–f**). Thus, readout alignment could be approximated from the local relationship between individual neuronal activity and the tendency to generate an action.

We next introduced a local plasticity rule that continuously updated thalamocortical feedback during task performance. Postsynaptic dendritic spikes induced by thalamocortical activity, together with the selected action, determined the targets of plasticity, whereas the RPE determined the direction of the change (**Fig. 6g**). Positive RPE induced long-term potentiation, whereas negative RPE induced long-term depression. Adding this dendritic-spike- and RPE-dependent thalamocortical plasticity increased the number of rewards, particularly during late learning, relative to the Full model with fixed readout alignment (**Fig. 6h**). This improvement was accompanied by a reduction in the number of invalid Actions (**Fig. 6i**). Thus, local calibration acquired readout alignment, whereas thalamocortical plasticity during task performance further refined behavioral output beyond that achieved with fixed alignment.

### The same closed-loop circuit supports single- and multi-target neural operant conditioning

BMI learning requires volitional control at the level of neuronal activity. We therefore asked whether module-specific parallel feedback was required in the BWSLC model to assign reinforcement-learning credit simultaneously to multiple cortical target cells during BMI learning. We used the same BWSLC circuit as in the cue-triggered action task, removed the Cue input, and generated reward directly from cortical neuronal activity (**Fig. 7**).

**Figure 7.**
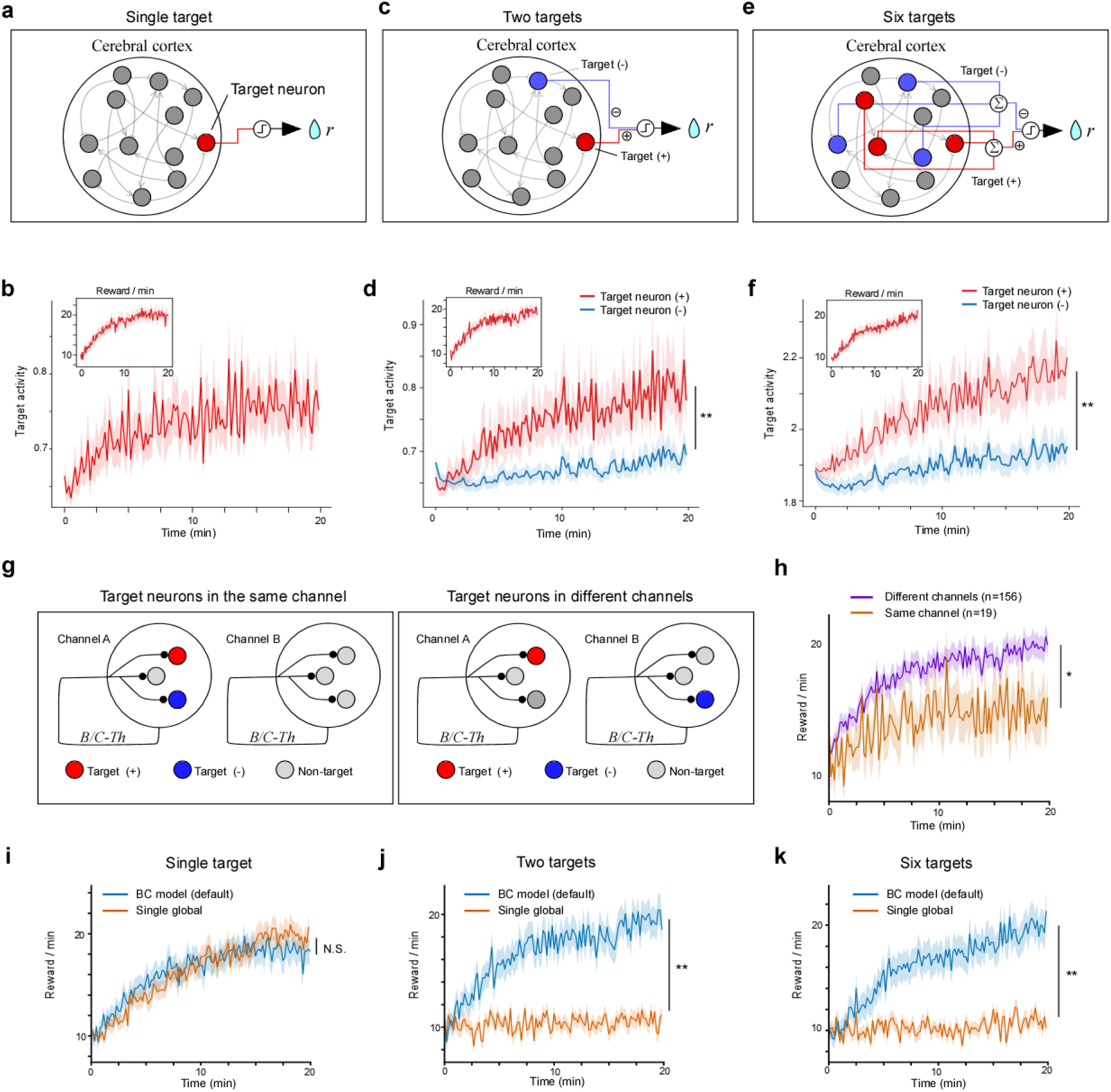
| Multi-module architecture enables multi-target neural operant conditioning. a, Schematic of the single-target condition. The activity of one cortical neuron was entered into the reward-generation function. **b,** Time course of target-neuron activity during learning. The inset shows the number of rewards per minute. **c,** Schematic of the two-target condition. The activity of the positive target neuron (red) increased reward, whereas the activity of the negative target neuron (blue) decreased reward. **d,** Time course of positive- and negative-target-neuron activity during learning. During the final 2 min of learning, positive-target-neuron activity was significantly higher than negative-target-neuron activity (Wilcoxon signed-rank test, *p=0.023, n= 100). **e,** Schematic of the six-target condition. The activities of three positive targets and three negative targets were integrated separately to calculate reward. **f,** Time course of positive- and negative-target-population activity during learning. The inset shows the number of rewards per minute. During the final 2 min of learning, summed positive-target activity was significantly higher than summed negative-target activity (Wilcoxon signed-rank test, **p=1.5 × 10^-7^, n=100 simulation sessions). **g,** Schematic of conditions in which the positive and negative target neurons belonged to the same channel or to different channels. **h,** Time course of the number of rewards per minute when targets were placed in different channels (purple) or in the same channel (orange). During the final 2 min of learning, reward rate was higher when the targets were placed in different channels than when they were placed in the same channel (Wilcoxon rank-sum test, *p=0.049, n=156 and 19 simulation sessions). **i,** Number of rewards in the standard BC model (blue) and a model using a single global feedback signal (orange) under the single-target condition. No significant difference was detected between the models (Wilcoxon signed-rank test, p=0.19, n=100). **j,** Number of rewards in the two models under the two-target condition. Performance was significantly higher in the BC model (Wilcoxon signed-rank test, **p=2.0×10^-6^, n=100). **k,** Number of rewards in the two models under the six-target condition. Performance was significantly higher in the BC model (Wilcoxon signed-rank test, **p=2.0×10^-8^, n=100).

In the single-target condition, reward was delivered when the activity of one cortical neuron increased (**Fig. 7a,b**). In the two-target condition, the activity of one positive target neuron increased reward, whereas the activity of a separate negative target neuron decreased reward (**Fig. 7c,d**). In the six-target condition, reward was calculated from the difference between the summed activity of three positive target neurons and that of three negative target neurons (**Fig. 7e,f**). In all conditions, the number of rewards increased with learning, and the activity of the target neuron or target population changed in the direction that increased reward. Similar circuit ablations, plasticity-off manipulations, and identity shuffling reduced learning performance in each condition (**Extended Data Fig. 4**).

In the two-target condition, we compared conditions in which the positive and negative target neurons belonged to the same module or to different modules (**Fig. 7g**). The number of rewards was higher when the two targets belonged to different modules than when they belonged to the same module (**Fig. 7h**). Thus, separating targets conditioned in opposing directions into distinct parallel modules improved learning performance.

Finally, we replaced module-specific thalamocortical feedback with a single global feedback signal. In the single-target condition, performance did not differ between the standard BWSLC model and the global-feedback model (**Fig. 7i**). In contrast, replacing module-specific feedback with global feedback markedly reduced the learning-related increase in reward number in the two-target and six-target conditions (**Fig. 7j,k**). Thus, module-specific feedback was dispensable when credit was assigned to a single target but was required when multiple cortical populations had to be modified simultaneously in different directions. These results indicate that parallel modules enable multi-target neural operant conditioning by receiving a common global reward while maintaining learning credit separately within each module.

### BWSLC architecture supports two-alternative forced choice

To determine whether the BWSLC model could be applied in a broader context, we trained it on a decision-making task in which it had to select the appropriate one of two actions in response to one of two Cue inputs (**Extended Data Fig. 5a**). The BWSLC model learned the task immediately (**Extended Data Fig. 5b**). In the circuit-manipulation experiments, all manipulations significantly affected performance except shuffling of the additive thalamocortical input (**Extended Data Fig. 5c**). The model behaved probabilistically in response to ambiguous stimuli and exhibited a smooth psychometric curve (**Extended Data Fig. 5d,e**). These results indicate that the BWSLC model is also applicable to tasks requiring a choice between two actions.

## Discussion

In this study, we showed that thalamic neurons receiving input from the SNr or LCN occupied largely distinct regions within the thalamus, with partial overlap in VM, while their cortical axons shared an L1a-enriched distribution, most prominently in M2. Related superficial projection components were also observed in M1 and S1, and BSVT analysis integrating publicly available anatomical datasets reproduced this upper-L1 return pattern, particularly in M2. In the BWSLC model incorporating this anatomical organization, reinforcement learning was most efficient when basal ganglia and cerebellar outputs returned to the cortex through both additive drive and gain modulation, and when this feedback was aligned with each cortical neuron’s contribution to behavioral output. A local activity-based rule for acquiring thalamocortical alignment and thalamocortical plasticity dependent on dendritic spikes and reward-prediction errors restored or refined learning. Furthermore, module-specific parallel feedback was required to condition the activities of multiple cortical neurons simultaneously in different directions. Together, these findings provide a framework in which basal ganglia– cerebellar cooperation is understood not as the simple summation of two outputs, but as a brain-wide credit-routing architecture comprising laminar reconvergence, thalamocortical alignment, local plasticity, and parallel modules.

The anatomical significance of our findings lies not in the individual observation that both basal ganglia and cerebellar pathways reach cortical L1, but in the reconvergence of two channels that remain largely segregated within the thalamus onto a shared laminar compartment in the cortex. This organization spans at least three spatial levels. First, SNr-recipient and LCN- recipient thalamic neurons are broadly separated at the level of thalamic nuclei and regions. Second, a subset of cells in VM receives input from both pathways, producing local mixing at the cellular level. Third, axons arising from otherwise distinct thalamic populations share a preferential distribution in cortical L1a, producing laminar reconvergence. The prominence of this shared L1a component in M2 suggests that higher-order motor cortex serves as a major interface through which largely segregated basal ganglia and cerebellar outputs are returned to a common cortical state. At the same time, each pathway retains distinct projection components in the middle and deep layers, indicating that the two systems do not become fully homogenized in the cortex but instead combine a shared L1 component with pathway-specific laminar components. This organization, in which segregation is preserved while the pathways reconverge, connects the classical view of the basal ganglia and cerebellum as independent loops with the more recent view of the two systems as components of an integrated network (Aoki et al., 2019; Bostan and Strick, 2018; Foster et al., 2021; Pisano et al., 2021).

Distal apical dendrites concentrated in L1 can impose context-dependent amplification, gating, and burst generation on cortical states already established by perisomatic and proximal inputs (Larkum et al., 1999, 2004; Xu et al., 2012). By converging in M2 L1a, the SNr-recipient and LCN-recipient pathways could therefore influence a common ongoing cortical state and selectively amplify it according to value, error, and behavioral context. In contrast, feedback components reaching the middle and deep layers of M2 can more directly drive local circuits and cortical output neurons. In the BWSLC model, these two effects were implemented as gain modulation and additive drive, respectively. The anatomy establishes laminar convergence; convergence onto individual cortical neurons, gain modulation, and readout alignment are hypotheses explored in the model. The finding that learning performance was highest when both components were present suggests that thalamocortical feedback can coordinate two complementary operations: selecting or amplifying a cortical state and directly driving that state toward action. Together with evidence that learning enhances behaviorally relevant representations in apical dendrites and that thalamic influence selectively reorganizes motor cortical populations after skill acquisition, our results position M2 L1a as an apical control interface through which action-related information evaluated in subcortical circuits can be incorporated into ongoing cortical population dynamics (Benezra et al., 2024; Cichon and Gan, 2015; Ramot et al., 2025).

For value-dependent changes generated in the basal ganglia and cerebellum to produce effective learning, the destination of the returning signal is as important as its magnitude. In the BWSLC model, aligning thalamocortical feedback with each cortical neuron’s contribution to action readout selectively returned learning credit to the populations most responsible for generating the action. The reduction in performance caused by shuffling feedback identity indicates that thalamocortical feedback acts as a routing signal that distributes credit across cortical populations. Moreover, high learning performance was restored by estimating each neuron’s readout contribution from the local relationship between its activity and the tendency to generate an action during spontaneous behavior, and then using this estimate to adjust thalamocortical weights. This local calibration indicates that a closed-loop circuit can use the correlation between neuronal activity and action probability to align forward contribution with return feedback. This principle is related both to node-perturbation learning, which estimates behavioral gradients from activity perturbations, and to feedback alignment, in which useful feedback directions emerge without an exact copy of the forward weights (Fiete and Seung, 2006; Fiete et al., 2007; Lillicrap et al., 2016). In the BWSLC, these general learning principles are assigned to the anatomically defined cortex–basal ganglia/cerebellum–thalamus–cortex return pathway. In addition, a three-factor plasticity rule combining presynaptic thalamocortical activity, postsynaptic dendritic spikes, and a global reward-prediction error provided a mechanism for experience-dependent refinement of routing alignment. The selective reduction in invalid actions during late learning indicates that thalamocortical plasticity not only supports the acquisition of new outputs but also removes unnecessary outputs from an acquired policy and refines the cortical-to-behavioral mapping (Biane et al., 2016; Frémaux and Gerstner, 2016; Gerstner et al., 2018; Tanaka et al., 2018).

The BWSLC extends the basal ganglia–cerebellar synergy described by Yoshida et al. (2025) into an explicit closed-loop credit-routing architecture. In the previous model, the basal ganglia direct pathway and the cerebellar parallel fiber–Purkinje cell pathway underwent plastic changes toward a common value objective, and learning was facilitated when their distinct temporal responses formed an appropriate phase relationship. The BWSLC retains this temporal cooperation while introducing a recurrent organization in which subcortical outputs return through the thalamus to a cortical recurrent neural network and thereby alter subsequent cortical states and action selection. Because cortical recurrent and readout weights remained fixed in the Full model, subcortical plasticity alone was sufficient to produce learned changes in cortical activity and behavior. The question of when the basal ganglia and cerebellum cooperate is therefore extended to the question of when, where, and in what form learning credit is returned. Within a single closed loop, the model integrates temporal alignment between distinct response kernels, spatial alignment with cortical neurons responsible for action readout, compartmental alignment through L1 and non-L1 pathways engaging different cortical computations, and module alignment that distributes credit across parallel loops. In contrast to the classical division of labor assigning reinforcement learning to the basal ganglia, supervised learning to the cerebellum, and representational learning to the cortex (Houk and Wise, 1995; Doya, 1999), our model shows that, during reward-contingent action learning, the two subcortical systems can share a common value objective while improving the same cortical policy through distinct temporal transformations and plasticity mechanisms. The finding that the LCN–thalamus–M2 pathway conveys reward-based post-error signals is also consistent with a framework in which the basal ganglia and cerebellum retain distinct internal computations while contributing to a shared behavioral objective (Ako et al., 2025). In this view, thalamocortical feedback is not merely the endpoint of two subcortical outputs, but a credit-routing interface that redistributes temporally processed subcortical signals to the appropriate cortical populations.

Module-specific parallel feedback was important for extending this credit-routing architecture to high-dimensional cortical control. Global feedback was sufficient when reward depended on the activity of a single target neuron, whereas module-specific feedback was required when two or six targets had to be adjusted simultaneously in opposing directions. Multi-target learning therefore requires more than broadcasting a single scalar reward throughout the network: conflicting updates must be separated into independent credit domains to prevent interference among local plasticity processes. The modules in the BWSLC can be understood not as fixed anatomical cortical boundaries, but as functional populations or feedback subspaces that share learning credit through a common basal ganglia–cerebellar–thalamic return pathway. This organization is related to mixture-of-experts and MOSAIC architectures, but differs in what is being routed. In mixture-of-experts models, an appropriate expert is selected according to its responsibility for a particular input, state, or environmental dynamics (Jacobs et al., 1991; Wolpert and Kawato, 1998; Haruno et al., 2001). In the BWSLC, by contrast, credit arising from the same global reward is routed to distinct cortical populations according to their responsibility for behavioral output. Thus, whereas mixture-of-experts architectures primarily route information or policies, the BWSLC routes learning credit. This principle also provides a circuit framework for understanding neural operant conditioning and BMI learning, in which the activity of small numbers of cortical neurons is brought under volitional control through reward (Fetz, 1969; Hira et al., 2014; Clancy et al., 2014). The requirement for the basal ganglia during BMI learning (Koralek et al., 2012; Neely et al., 2018), together with the faster acquisition of decoders aligned with pre-existing population manifolds (Sadtler et al., 2014; Athalye et al., 2017), suggests that cortical learnability may be constrained not only by intracortical dynamics but also by alignment with pre-existing loop-specific feedback. The BWSLC therefore recasts the natural manifold as both a subspace of cortical activity and a credit-routing subspace that is preferentially accessible to subcortical learning signals. In this framework, partially segregated basal ganglia and cerebellar channels reconverge through an L1a-enriched thalamocortical interface, most prominently in M2, and use plastic, module-specific feedback to route reinforcement-learning credit to the cortical populations responsible for action.

## Methods

### Animals

Animal procedures for the viral pathway-tracing experiments were approved by the Animal Research Ethics Committee of the Institutional Animal Care and Use Committee of the Institute of Science Tokyo (A2021-290C6, A2024-060C2). Animal procedures for the M2 tracer injection experiments were approved by the Institutional Animal Care and Use Committee of the National Institutes of Natural Sciences, Japan. All experiments were conducted in accordance with the Basic Guidelines for the Conduct of Animal Experiments and Related Activities in Research Institutions issued by the Ministry of Education, Culture, Sports, Science and Technology of Japan. A total of 12 adult C57BL/6 mice (6 males and 6 females) and 2 Ai162 transgenic mice (Daigle et al., 2018) that can express GCaMP6s in a Cre-dependent manner were used for the experiments. Mice were 6–24 weeks old at the time of injection.

### Stereotaxic viral injections

Mice were anesthetized with 1.5% isoflurane and secured in a stereotaxic frame. Anesthesia was maintained using air or oxygen-enriched air delivered at 0.8–1.0 l min^−1^. Body temperature was maintained throughout surgery using a heating pad. The scalp was incised, connective tissue was removed from the skull, and a small craniotomy was made above each target site. Viruses were pressure-injected through borosilicate glass pipettes. Injection volumes were 200 nl for the SNr, 500 nl for the LCN or MCN, and 500 nl for the thalamus. Viruses were injected at 50 nl min^−1^. After each injection, the pipette was left in place for 10 min to minimize backflow before being slowly withdrawn. After surgery, mice were allowed to recover on a heating pad before being returned to their home cages. To label pathways originating from the substantia nigra pars reticulata (SNr), AAV1-EF1a-FlpO (1.15 × 10^13^ gc ml^−1^) or AAV1-hSyn-Cre-WPRE (1.2 × 10^13^ gc ml^−1^) was injected into the SNr ipsilateral to the thalamic injection site. To label pathways originating from the cerebellar nuclei, the corresponding vector was injected into the lateral cerebellar nucleus (LCN) or medial cerebellar nucleus (MCN) contralateral to the thalamic injection site. FlpO- or Cre-dependent fluorescent reporter AAVs, AAV-CAG-FRT-Rev-tdTomato (5.7 × 10^12^ gc ml^−1^) or AAV-pCAG-FLEX-EGFP-WPRE (6.25 × 10^12^ gc ml^−1^), respectively, were injected into the thalamus to label thalamic neurons receiving input from the injected SNr or cerebellar nucleus.

To examine double labeling in the thalamus, we used two Ai162 mice (Cre-dependent GCaMP6s reporter mice). AAV-CAG-FRT-Rev-tdTomato (5.7 × 10^12^ gc ml^−1^) was injected into the thalamus, AAV1-EF1a-FlpO (1.15 × 10^13^ gc ml^−1^) into the SNr, and AAV1-hSyn-Cre-WPRE (1.2 × 10^13^ gc ml^−1^) into the LCN.

To determine the local extent of viral transduction, AAV5-CAG-tdTomato (1.0 × 10^11^ gc ml^−1^), AAV5- CAG-GFP (1.0 × 10^11^ gc ml^−1^), or a mixture of AAV-EF1a-FlpO (2.0 × 10^12^ gc ml^−1^) and AAV-CAG-FRT-Rev-tdTomato (1.71 × 10^13^ gc ml^−1^) was injected into the SNr, LCN, MCN, or thalamus.

To retrogradely label subcortical neurons projecting to the thalamus, AAV2retro-CAG-GFP (5.0 × 10^12^ gc ml^−1^) was injected into the thalamus (Tervo et al., 2016), and retrogradely labeled neurons were subsequently examined in the SNr and cerebellar nuclei.

Stereotaxic coordinates were defined relative to bregma and varied slightly across animals as follows: thalamus (VM region), anteroposterior (AP) −1.5 to −1.6 mm, mediolateral (ML) 0.9 mm, and dorsoventral (DV) −3.5 to −4.0 mm; SNr, AP −3.0 to −3.3 mm, ML 1.5 to 1.6 mm, and DV −4.3 to −4.9 mm; LCN, AP −6.1 to −6.2 mm, ML 1.8 to 2.1 mm, and DV −2.1 to −2.2 mm; and MCN, AP −6.3 mm, ML 0.7 mm, and DV −2.1 to −2.2 mm.

### Histological processing for viral pathway-tracing experiments

Three weeks after viral injection, mice were deeply anesthetized with isoflurane and transcardially perfused with phosphate-buffered saline (PBS), followed by 4% paraformaldehyde (PFA). Brains were removed and postfixed in 4% PFA at 4°C for 2 days and then immersed in 30% sucrose in PBS for several days until they sank. Brains were rapidly frozen in isopentane cooled on dry ice and stored at −80°C until sectioning. Sagittal sections were cut at a thickness of 20 μm at −20°C using a cryostat (NX50, Epredia). Sections were collected across levels containing the cerebral cortex, thalamus, SNr, and cerebellar nuclei. Sections were mounted on glass slides, counterstained with DAPI, and coverslipped.

### Tracer injection and immunohistochemistry

Two male C57BL/6 mice aged 6 weeks were anesthetized with a mixture of ketamine (40 mg kg−1, i.p.) and xylazine (4 mg kg−1, i.p.) and placed in a stereotaxic apparatus. A mixture of anterograde and retrograde tracers was injected into secondary motor cortex (M2) at 2.5–2.8 mm anterior and 0.8 mm lateral to bregma and 0.2–0.8 mm below the pial surface. Glass pipettes were filled with 0.5 M potassium acetate containing 2.5% (w/v) biotinylated dextran amine (BDA; BDA-10000, Invitrogen) and 0.2% (w/v) Alexa Fluor 555-conjugated cholera toxin subunit B (CTB555; Invitrogen). A total volume of 70 nl was pressure-injected into M2 using a PV820 pressure injector (World Precision Instruments). After a survival period of 7 days, animals were deeply anesthetized with sodium pentobarbital (60 mg kg−1, i.p.) and transcardially perfused with 4 ml of a prefixative solution containing 250 mM sucrose and 5 mM MgCl2 in 0.02 M phosphate buffer (PB; pH 7.4), followed by a fixative solution containing 4% paraformaldehyde and 0.2% picric acid in 0.1 M PB. Brains were cut into 50-μm sagittal sections using a vibratome (Leica). For visualization of BDA, sections were incubated overnight in a 1:100 dilution of avidin-biotinylated horseradish peroxidase complex (ABC Elite; Vector Laboratories) in 0.05 M Tris-HCl-buffered saline (TBS). The BDA signal was enhanced by incubation for 30 min with 2.5 μM biotinylated tyramine, 3 μg ml−1 glucose oxidase, and 2 mg ml−1 β-D-glucose. Sections were subsequently incubated with Alexa Fluor 488-conjugated streptavidin (Invitrogen).

For immunofluorescence staining, sections were incubated with a guinea pig polyclonal antibody against vesicular glutamate transporter type 2 (VGLUT2; AB2251, EMD Millipore; 1:500), a mouse monoclonal antibody against neurofilament heavy chain (NF-H; N-200, N0142, Sigma-Aldrich; 1:1000), a mouse monoclonal antibody against neuronal nuclei (NeuN; MAB377, EMD Millipore; 1:2000), a rabbit polyclonal antibody against calbindin D-28K (CB-38a, Swant; 1:1000), and a mouse monoclonal antibody against glutamate decarboxylase 67 (GAD67; MAB5406, EMD Millipore; 1:1000) in 0.05 M TBS containing 10% normal goat serum, 2% bovine serum albumin, and 0.5% Triton X-100. After washing with TBS, sections were incubated with appropriate Alexa Fluor-conjugated secondary antibodies (1:200; Life Technologies) for 2–3 h at room temperature. Sections were mounted on glass slides using Vectashield mounting medium (Vector Laboratories). The borders between M2 and primary motor cortex (M1) were identified using NF-H (N-200) immunoreactivity.

### Fluorescence imaging

Fluorescence images were acquired using a BZ-X700 microscope (Keyence) equipped with a 20× objective lens. For each animal, sections containing the injection sites, labeled cells in the thalamus, and axonal projections in the cerebral cortex were imaged. When a region of interest could not be captured within a single field of view, multiple fields were acquired. Images were saved in TIFF format and analyzed using ImageJ, MATLAB, or Python. Thalamic nuclei were identified by overlaying nuclear boundaries from the corresponding atlas sections onto the histological sections. For the quantification in Fig. 2g,h, a total of 2,108 labeled thalamic cells across 42 unique section records from three mice in the SNr group and 1,174 cells across 39 unique section records from three mice in the LCN group were identified and counted.

### Quantification of thalamocortical axon distributions

To assess the laminar distribution of thalamocortical axons, fluorescence intensity was first quantified along cortical depth. In each M1, M2, and S1 image, a region of interest extending from the pial surface to the white-matter boundary was defined. Mean fluorescence intensity was calculated within successive depth bins, and background fluorescence was subtracted. Background values were estimated either from an adjacent region lacking labeled axons or from the distribution of low-intensity pixels within the image. Fluorescence-intensity profiles were plotted against cortical depth and aligned with cortical layer and areal boundaries identified from DAPI images. This analysis was used to determine the cortical layers in which axons of SNr-, LCN-, or MCN-recipient thalamic neurons were preferentially distributed in M1, M2, and S1. Because fluorescence intensity may underestimate dim axons or thin, spatially dispersed axonal structures, total axon length was also quantified within each cortical layer. After background subtraction, candidate axons were extracted by thresholding. Cell bodies, blood vessels, debris, saturated signals, and non-axonal signals arising from the injection site were excluded on the basis of size and shape and by visual inspection. The resulting axon masks were skeletonized to generate one-pixel-wide representations of the labeled axons. Total skeleton length was calculated separately for each cortical layer. This analysis was performed in M1, M2, and S1 for thalamocortical pathways defined by input from the SNr, LCN, or MCN.

### Statistical analysis of anatomical data

Pairwise comparisons were performed using the Wilcoxon rank-sum test for independent samples or the Wilcoxon signed-rank test for paired samples. Nucleus-wise cell-count and cell-density profiles were compared between the SNr and LCN groups using studentized Freedman–Lane permutation tests with 100,000 within-animal residual permutations. Holm-adjusted *P* values were reported where correction for multiple comparisons was applied. Projection patterns across the five most densely innervated thalamic nuclei were compared using exact PERMANOVA with all possible group-label assignments. CN- and SNr-weighted layer fractions from each source experiment were CLR-transformed and compared by exact PERMANOVA, with Holm correction across three areas and two tracing methods. Inference was conditional on the estimated thalamocortical output profiles. The equal-nucleus BSVT/BBVT comparison with the fixed VGLUT2 profile was descriptive. Separate source-balanced comparisons used sign-flip tests and were not used to assign significance to the equal-nucleus curves. Statistical analyses were performed using MATLAB R2024a (MathWorks) or Python 3.10. Data were presented as mean ± standard deviation (SD) unless otherwise indicated. Sample sizes were not determined by an a priori power calculation. Animals were excluded when viral expression was absent or when the injection site was substantially displaced from the intended target nucleus. For SNOC target and parallel/global comparisons, the five target-set sessions were averaged within each seed before paired testing.

### Virtual tracing

We integrated source-to-thalamus input distributions from a cerebellar nuclear group not classified as inhibitory-specific (CN) or the substantia nigra pars reticulata (SNr) with nucleus-specific thalamocortical output distributions to estimate relative cortical-layer profiles for CN→thalamus→cortex and SNr→thalamus→cortex pathways. Thalamocortical outputs were quantified independently from BIL/BICCN single-neuron reconstructions and bulk anterograde tracing in the Allen Mouse Brain Connectivity Atlas. CN comprised six non-inhibitory-specific experiments in which the dentate nucleus was a primary or secondary injection structure; seven SNr experiments were retained. For each source group, projection density in each thalamic nucleus was divided by source injection volume and averaged across experiments:

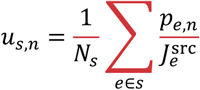

Cortical layers were defined using the Allen Mouse Brain Common Coordinate Framework (CCF) 100- µm annotation (Wang et al., 2020a) and ccf-streamlines. Each cortical voxel was assigned to its nearest streamline, and relative depth within the parent layer was calculated as:

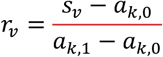

For L1 and L5, depths below 0.5 were classified as upper and those at or above 0.5 as lower. Allen labels for L2/3, L4, L6a, and L6b were retained. Voxels lacking a valid streamline lookup used the previous distance-based assignment. The same mask was applied to single-neuron and bulk datasets. We analyzed neuronal-morphology reconstructions in SWC format from 195 BIL/BICCN thalamic neurons. Coordinates were interpreted in CCF micrometers, type-2 segments were treated as axons, and segments were resampled every 10 µm. Axon length was accumulated for each cortical area and layer, divided by area-layer volume, and averaged among neurons with somata in the same thalamic nucleus. Missing cell-area-layer combinations were set to zero before averaging. A representative VM neuron was selected by ranking the number of M1/M2/S1 targets containing L1 axon and then total L1 axon length.

For bulk outputs, 103 experiments with primary thalamic injection sites were analyzed. Projection density was averaged within each area-layer mask, divided by injection volume, and then averaged across experiments sharing the same primary nucleus. The archived analysis averaged all voxels in each anatomical mask without applying the Allen data_mask; a valid-voxel sensitivity analysis is required to assess missing-data effects.

For source s and nucleus n, the source-to-thalamus weight was the product of estimated neuron number and mean normalized source input density. This relative weight was not interpreted as an absolute synapse count. The primary comparison used 12 nuclei represented in both output methods (Allen ontology acronyms): CM, IMD, PIL, PO, PVT, RE, RT, SMT, VAL, VM, VPL, and VPMpc. Source-specific cortical output was estimated by summing nucleus-specific output densities weighted by source-to-thalamus input: Estimated neuron numbers were obtained by multiplying each nucleus volume in the 25-µm CCF annotation by the corresponding Blue Brain Cell Atlas neuron-density estimate; the region matches and values are provided with the code.

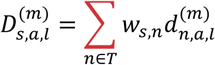

M1 was defined as MOp, M2 as MOs, and S1 as the volume-weighted aggregate of seven SSp subdivisions. Layer fractions were normalized within each method, source, and cortical area:

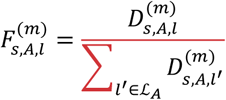

For comparison with VGLUT2 (Extended Data Fig. 2d), the equal-nucleus control used the arithmetic mean of thalamocortical output densities across the 12 matched nuclei, assigning each nucleus a weight of 1/12. The same cortical-area aggregation and layer normalization were then applied.

Analyses used Python 3.10, NumPy, pandas, SciPy, Matplotlib, AllenSDK, h5py, pynrrd, and ccf-streamlines.

### Cue-triggered action task

Following Yoshida et al. (2025), the default model was a basal ganglia–cerebellum model in which plasticity was restricted to selected basal ganglia and cerebellar pathways, corresponding to the Full model (BC in the figures; syn-BC in the code). The recurrent neural network (RNN) representing the cerebral cortex consisted of eight parallel channels, or modules, with 25 units per module and 200 units in total. Each module contained 10 excitatory Context units, five excitatory Action units, five excitatory No-Action units, and five inhibitory units. Neural activity was updated at 1-ms resolution with a time constant of 20 ms. The RNN had sparse recurrent connectivity that obeyed Dale’s law. Recurrent weights remained fixed throughout all tasks.

Activity of the Context units in each module was read out through six pathways. The basal ganglia pathways comprised the hyperdirect (HD), direct (D), and indirect (ID) pathways, whereas the cerebellar pathways comprised P1, P2, and P3. Integrated basal ganglia and cerebellar outputs were defined as

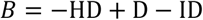

and

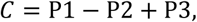

respectively. The resulting basal ganglia and cerebellar signals were returned to the cortical RNN.

The pathway-specific delays before feedback reached the cortex were 29, 36, 49, 40, 40, and 80 ms for HD, D, ID, P1, P2, and P3, respectively. Positive integrated feedback additively drove the Action population, whereas negative integrated feedback additively drove the No-Action population. When the basal ganglia and cerebellar outputs had the same sign, an additional multiplicative gain of 1–3 was applied. The cell-specific feedback weights were set equal to the corresponding behavioral readout weights. Thus, the feedback pathway was structurally aligned with motor output from model initialization.

Tone input was delivered to the Context and inhibitory units. The intertrial interval was 1,000 ms after the end of the preceding action. Actions occurring 100–1,000 ms after tone onset were eligible for reward.

At each 1-ms time step, the activities of the Action and No-Action populations were aggregated using fixed readout weights. A slowly varying baseline was subtracted to obtain the action-value-like signals

*q*_action_ and *q*_no_. Within each 100-ms block, the time point at which *q*_action_ − *q*_no_ reached its maximum was selected. The difference between the two values at the corresponding RNN state was transformed through a logistic function to obtain the probability of an action. The final binary action was determined by stochastic sampling from this probability.

Learning was driven by a temporal-difference reward-prediction error (TD-RPE). The RPE was calculated from the current reward, the cost of the selected action, and the current and subsequent cortical action values:

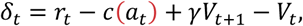

where *δ_t_* denotes the RPE, *r_t_* the reward, *c*(*a_t_*) the cost of the selected action, *V_t_* and *V_t_*_+1_ the values obtained from cortical readouts at the current and subsequent time points, and *γ* the temporal discount factor.

Plasticity was restricted to the basal ganglia direct pathway (D) and the parallel-fiber input to Purkinje cells (P2). Context activity during the immediately preceding 100-ms block, after subtraction of a slow baseline, was used to define the eligibility trace. The eligibility trace was multiplied by the RPE, and weights were updated every 1 s. A positive learning signal induced long-term potentiation (LTP) in D and long-term depression (LTD) in P2. A negative learning signal induced LTD in D and LTP in P2. After each update, the total weight within each module was renormalized to its initial value.

Control manipulations included a linear-only condition, in which only additive feedback was retained; a gain-only condition, in which only multiplicative gain was retained; and a no-feedback condition, in which both feedback components were removed. Plasticity was also disabled selectively in D or P2, and basal ganglia or cerebellar nuclear output was suppressed. In the identity-shuffle conditions, the magnitude and temporal waveform of each feedback signal were preserved, whereas the cell-specific profile of its target weights was randomized. This manipulation tested whether the correct correspondence between feedback identity and cortical target cells was required.

### Thalamocortical plasticity

In the Full model, the cell-specific profile of thalamocortical feedback was fixed. In the BC+TCP model, thalamocortical plasticity (TCP) was added, allowing the cell-specific gain of the additive current delivered by positive basal ganglia or cerebellar feedback to Action units to be modified by learning.

Thalamocortical eligibility was calculated as the product of positive basal ganglia or cerebellar feedback that had reached the cortex during the preceding block, positive phasic activity of the Action unit, the fixed feedback/readout profile, a local gain gate, and a postsynaptic activity gate. The postsynaptic activity gate served as a proxy for dendritic-spike-dependent plasticity. Thus, credit was assigned to a thalamocortical connection only when subcortical input reached the cortex and an Action unit aligned with the direction of that input was sufficiently active.

For actions accompanied by a positive RPE, the thalamocortical gain from the basal ganglia or cerebellar pathway to the Action unit underwent LTP. For unrewarded actions outside the reward window, termed invalid actions, a negative RPE induced LTD of the same thalamocortical gain. Gains were initialized at 1.0 and constrained to the range 0–2.5. The learning rates for LTP and LTD were 0.001125 and 0.00825, respectively, and weights were updated every 1 s. The weak positive-feedback-to-No-Action branch was initialized at 0.02.

### Single-neuron operant conditioning

For single-neuron operant conditioning (SNOC), the Full-model circuit was used with cue input removed and reward determined by target-neuron activity. Networks were independently initialized for each of 20 seeds without prior task training; five target-set sessions shared the same initial network within each seed. In this study, SNOC included a conventional single-target condition as well as two-target and six-target differential conditions.

Each session comprised a 2-min baseline period with no tone and no externally defined task reward, followed immediately by 20 min of SNOC. D and P2 plasticity was disabled during baseline. A session-specific reward threshold was calculated from a target signal constructed from the maximum activity of each target neuron within each 100-ms bin.

In the single-target condition, the signal was the binwise maximum activity of one target neuron. In the two-target condition, the signal was the activity of target neuron 1 minus that of target neuron 2. In the six-target condition, the signal was the summed activity of three target neurons minus the summed activity of three other target neurons. For each session, the threshold was selected from baseline spontaneous activity to produce approximately 10 reward events per minute after accounting for a 1-s reward-refractory period.

During SNOC, reward was delivered when the target signal exceeded the session-specific threshold and at least 1 s had elapsed since the previous reward. Plasticity followed the same D/P2 learning rule as in the cue-triggered action task.

### Two-alternative forced-choice task

The basic Full-model architecture was retained for the two-alternative forced-choice task, with modifications to accommodate two inputs and two behavioral outputs. Action units in modules 0–3 were assigned to choice L (action 1), whereas Action units in modules 4–7 were assigned to choice R (action 2). The former Action populations therefore represented choices L and R, and the former No-Action populations were redefined as No-Go populations.

Cue L (Cue 1) and Cue R (Cue 2) were generated as independent signed random input vectors and delivered to the Context and inhibitory units in all eight modules. Intermediate cues were generated as weighted averages of Cue L and Cue R, with the two weights constrained to sum to one. Cue presentation began 50 ms after trial onset and continued until a choice was made or the trial timed out. Choices were permitted from 200 to 1,100 ms after trial onset.

Within each module, the No-Go readout was subtracted from the corresponding choice readout. Choice evidence was then summed across the four modules assigned to choice L and separately across the four modules assigned to choice R. Each summed signal was transformed through a logistic function to obtain an independent choice probability. Choices L and R were mutually exclusive. If both or neither choice was sampled, the outcome was treated as no-choice and the trial continued.

For each decision attempt, an action-specific policy score was calculated as the difference between the Bernoulli Go sample for that choice and its predicted choice probability:

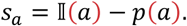

This score was multiplied by the Context eligibility traces in the four modules corresponding to that choice and accumulated until the end of the trial. A terminal TD-RPE was calculated after a correct choice, an incorrect choice, or a timeout. The terminal TD-RPE was multiplied by the accumulated trial-level eligibility, and the same opponent D/P2 update used in the cue-triggered action task was applied. The model was trained for 20 min using only the two endpoint cues, Cue L and Cue R. After training, D/P2 weight updates and weight noise were disabled. A 13.3-min psychometric test was then performed using 11 mixtures of Cue L and Cue R. A logistic model was fitted to the resulting psychometric function. The primary analysis was performed using 100 independent random seeds.

## Acknowledgements

We thank Re. Hira, T. Shimada, T. Nagai and all other members of the Isomura lab for supporting this work. This work was supported by JP22wm0525007 (RH), JP19dm0207089 (YI) from AMED, JP22H02731 (RH), JP20K22678 (RH), JP21H05134 (RH), JP21H05135 (RH), JP26H01161 (RH), JP26K23436 (RH), JP21H05242 (YI), and JP23H02589 (YI) from MEXT/JSPS, JPMJCR1751 (YI) from JST, Nakatani Foundation (RH), Shimadzu Foundation (RH), Takeda Science Foundation (RH), The Precise Measurement Technology Promotion Foundation (RH), Tateishi Science and Technology Foundation (RH), Research Foundation for OptoScience and Technology (RH), and NINS Grant for Young Researcher Collaboration (RH, MM). This work utilized the TSUBAME4.0 supercomputer at the Institute of Science Tokyo for analysis. We used ChatGPT-5 and ChatGPT-6 (OpenAI) for English–Japanese translation and data analysis during manuscript preparation. For data-analysis tasks, all AI-generated code was thoroughly checked and validated for reliability before use.

## Author contributions

S.T., M.M., K.F. conducted experiments and analyzed the data. R.H. analyzed data and conducted modeling and simulation. K.K., S.K. provided AAV vectors. Y.I., Y.K. supervised the projects. R.H. wrote the paper with input from all authors.

## Data availability

All data including microscopy images and segmentation results will be deposited in Figshare upon acceptance.

## Code availability

The simulation code and virtual-tracing code will be shared at GitHub upon acceptance.

---

**Extended Data Figure 1.**
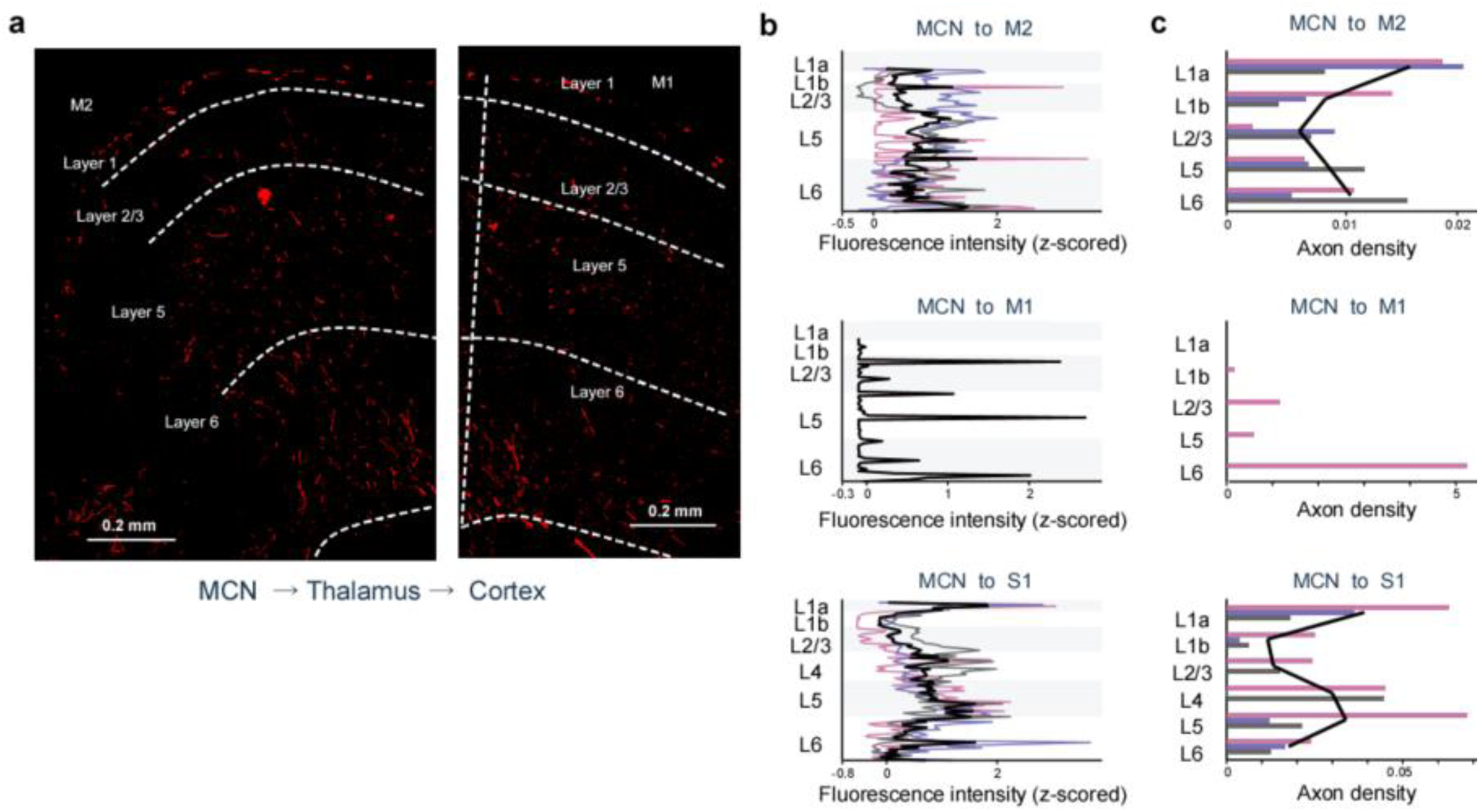
| Laminar distribution of the MCN-recipient thalamocortical pathway a, Representative labeled axons in M2 and M1 following transsynaptic labeling of the MCN-recipient thalamic pathway. Red, labeled axons; dashed lines, cortical layer boundaries. **b,** Background-subtracted, z-scored fluorescence profiles across cortical depth in M1, M2, and S1. Three mice were examined for each cortical area, but detectable labeled axons were found in M1 in only one of the three mice; the M1 profile therefore represents this single case. M2 and S1 profiles were obtained from all three mice. Colored lines represent individual mice, and black lines indicate the mean for M2 and S1. Labeled axons in M1 were sparse, whereas M2 and S1 showed both superficial and deep projection components. **c,** Layer-wise axon-length measurements in each cortical area. Bars represent individual mice, and black lines indicate the mean for M2 and S1.

**Extended Data Figure 2.**
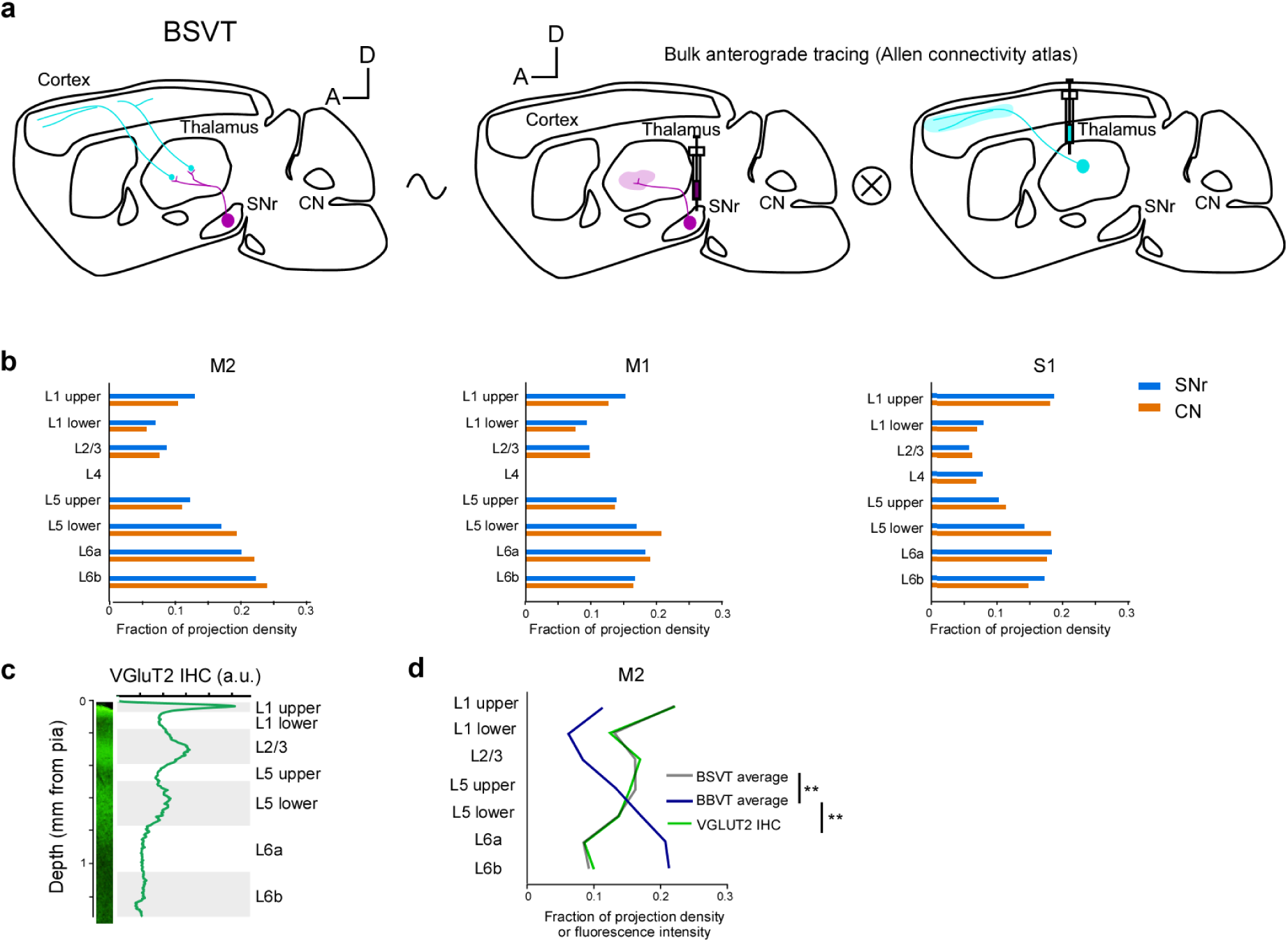
| BBVT and comparison with the VGLUT2 laminar profile **a,** Overview of Bulk-Bulk Virtual Tracing (BBVT), in which both source-to-thalamus and thalamus-to- cortex projections were estimated from bulk Allen connectivity experiments. **b,** BBVT estimates of relative layer fractions in M2, M1, and S1. **c,** Background-subtracted VGLUT2 fluorescence profile across cortical depth in the sagittal section used for the comparison. **d,** Layer-wise comparison of BSVT (gray), BBVT (blue), and normalized VGLUT2 fluorescence (green). BSVT and VGLUT2 retained larger superficial components, whereas BBVT was biased toward deep layers and differed significantly from both BSVT (CLR sign-flip tests, *p* = 2.4×10^-4^) and VGLUT2 (CLR sign-flip tests, *p* = 2.4×10^-4^). Both tracing estimates used the equal-weight mean across the 12 matched thalamic nuclei.

**Extended Data Figure 3.**
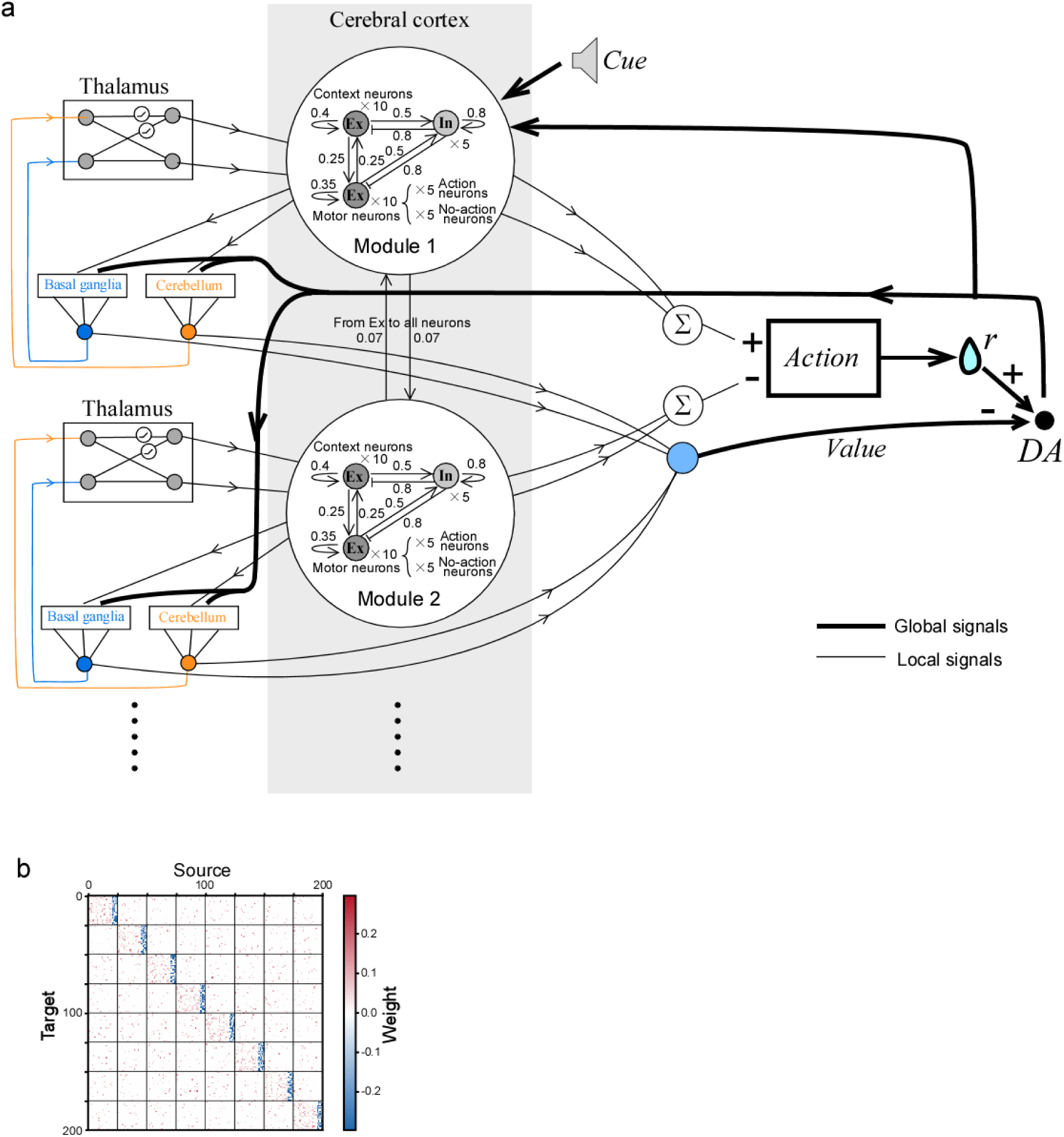
| Modular organization and recurrent connectivity of the cortical RNN a, Schematic of the BC model emphasizing its organization into eight parallel channels. Each module contained 10 Context neurons, 10 motor neurons consisting of five Action neurons and five No-Action neurons, and five inhibitory neurons. Feedback from the thalamus, basal ganglia, and cerebellum was provided in parallel to each module. Values shown on the connections indicate fixed connection weights or connection probabilities. The outputs of all channels were integrated to select an action, and the dopamine signal was calculated from the difference between reward r and the estimated value. **b,** Synaptic weight matrix of the cortical RNN, which consisted of eight modules and 200 neurons in total.

**Extended Data Figure 4.**
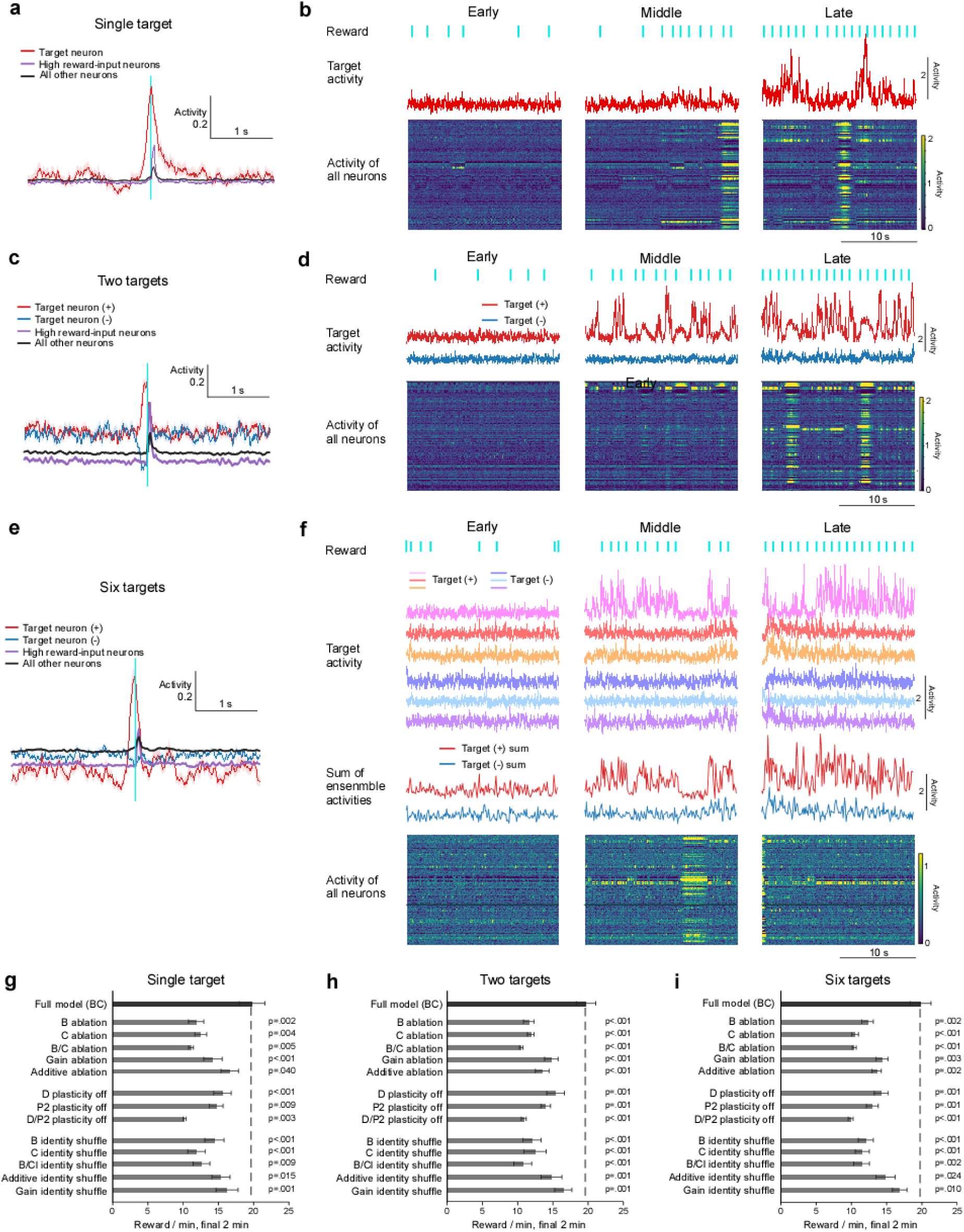
| Neural operant conditioning dynamics and circuit controls **a,** Mean activity of the reward-aligned target neuron (red), neurons receiving strong reward input (purple), and all other neurons (black). The cyan line indicates the time of reward delivery. **b,** Activity of the single target neuron and all neurons during early, middle, and late learning. Columns show successive learning stages. Cyan vertical lines indicate reward times. Traces show target- neuron activity, and heat maps show the activity of all neurons. **c,d,** Data corresponding to **a** and **b** for the two-target task. **e,f,** Data corresponding to a and **b** for the six-target task. Panel f additionally shows individual positive and negative targets and the summed activity of each target population. **g,** Number of rewards during the final 2 min under the Full model, signal-ablation, plasticity-off, and input-identity-shuffle conditions in the single-target condition. **h,** Number of rewards during the final 2 min after each model manipulation in the two-target condition. **i,** Number of rewards during the final 2 min after each model manipulation in the six-target condition. Mean ± SEM across 20 iterations. P values were obtained by comparison with the Full model using the Wilcoxon signed-rank test.

**Extended Data Figure 5.**
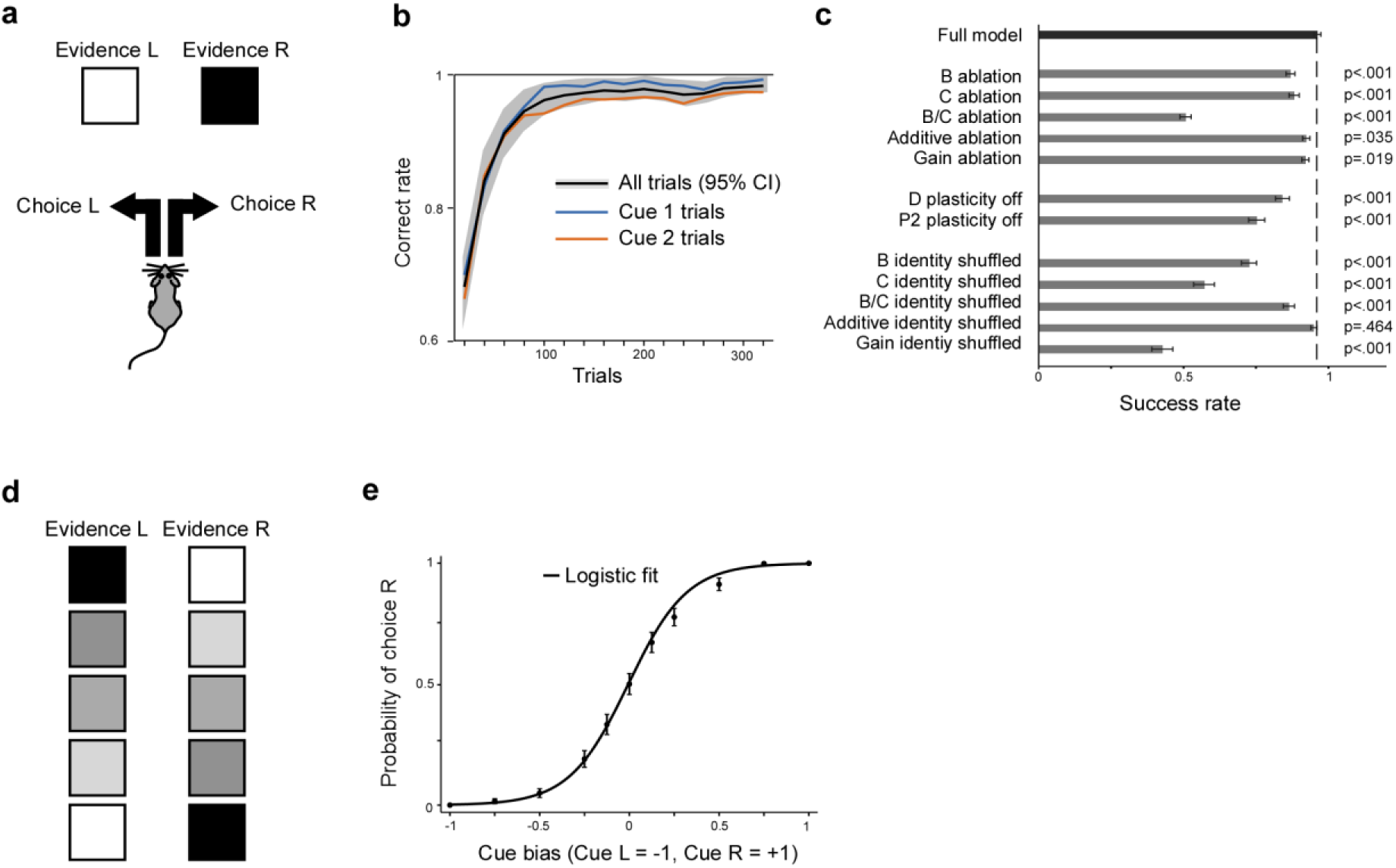
| A two-alternative forced-choice task and psychometric function **a,** Two-choice decision-making task. One of two Cues was presented, and the model selected one of two choices. A correct choice was rewarded. **b,** Learning curve for the task shown in **a**. **c,** Results of the circuit-manipulation experiments in the task shown in **a**. **d,** After learning the task shown in a, the two Cues were mixed and presented simultaneously. **e,** Psychometric function obtained in the experiment shown in **d**.

**Extended Data Table 1.** |Abbreviations of thalamic nuclei and neighboring regions Names for atlas-derived labels follow the Allen ontology.

| Abbreviation | Full name |
| --- | --- |
| AV | Anteroventral nucleus of thalamus |
| CL | Central lateral nucleus of the thalamus |
| CM | Central medial nucleus of the thalamus |
| Eth | Ethmoid nucleus of the thalamus |
| IGL | Intergeniculate leaflet of the lateral geniculate complex |
| IMD | Intermediodorsal nucleus of the thalamus |
| IntG | Intermediate geniculate nucleus |
| LD | Lateral dorsal nucleus of thalamus |
| LH | Lateral habenula |
| LP | Lateral posterior nucleus of the thalamus |
| MD | Mediodorsal nucleus of thalamus |
| MDm | Mediodorsal nucleus of the thalamus, medial part |
| MH | Medial habenula |
| PCN | Paracentral nucleus |
| PF | Parafascicular nucleus |
| PIL | Posterior intralaminar thalamic nucleus |
| POL | Posterior limiting nucleus of the thalamus |
| PO | Posterior complex of the thalamus |
| PoT | Posterior triangular thalamic nucleus |
| PP | Peripeduncular nucleus |
| PR | Perireunensis nucleus |
| PVT | Paraventricular nucleus of the thalamus |
| RE | Nucleus of reuniens |
| REth | Retroethmoid nucleus |
| RH | Rhomboid nucleus |
| RT | Reticular nucleus of the thalamus |
| SMT | Submedial nucleus of the thalamus |
| SPA | Subparafascicular area |
| SPFm | Subparafascicular nucleus, magnocellular part |
| SPFp | Subparafascicular nucleus, parvicellular part |
| SubG | Subgeniculate nucleus |
| VA | Ventral anterior thalamic nucleus |
| VAL | Ventral anterior-lateral complex of the thalamus |
| VL | Ventrolateral thalamic nucleus |
| VM | Ventral medial nucleus of the thalamus |
| VPL | Ventral posterolateral nucleus of the thalamus |
| VPLpc | Ventral posterolateral nucleus of the thalamus, parvicellular part |
| VPM | Ventral posteromedial nucleus of the thalamus |
| VPMpc | Ventral posteromedial nucleus of the thalamus, parvicellular part |
| Xi | Xiphoid thalamic nucleus |
| ZI | Zona incerta |

