## Supplementary information for "Thalamocortical credit routing from the basal ganglia and cerebellum: anatomical constraints and circuit principles"

**Affiliations:**
1. Department of Physiology and Cell Biology, Graduate School of Medical and Dental Sciences, Institute of Science Tokyo, 1-5-45 Yushima, Bunkyo-ku, Tokyo, 113-8510, Japan.
2. Graduate School of Brain Science, Doshisha University, Kyoto, Japan
3. Division of Cerebral Circuitry, National Institute for Physiological Sciences, Okazaki, Japan
4. Faculty of Pharmacy, Musashino University, Tokyo, Japan
5. Department of Molecular Genetics, Institute of Biomedical Sciences, Fukushima Medical University, Fukushima, 960-1295, Japan
6. Brain Science Institute, Tamagawa University, Tokyo, Japan
7. High Performance Artificial Intelligence Systems Research Team, Center for Computational Science, RIKEN, Tokyo 103-0027, Japan

[†] Corresponding authors:
Riichiro Hira, M.D., Ph.D.
Department of Physiology and Cell Biology, Graduate School of Medical and Dental Sciences, Institute of Science Tokyo, 1-5-45 Yushima, Bunkyo-ku, Tokyo, 113-8510, Japan.
Yoshikazu Isomura, Ph.D.
Department of Physiology and Cell Biology, Graduate School of Medical and Dental Sciences, Institute of Science Tokyo, 1-5-45 Yushima, Bunkyo-ku, Tokyo, 113-8510, Japan.

Contents

1. Cue-triggered action task and shared BWSLC implementation
   1.1 Cortical RNN architecture and activity dynamics
   1.2 Recurrent connectivity
   1.3 Context activity and six corticofugal pathways
   1.4 BG/CB integration, thalamocortical return, and cortical feedback
   1.5 Alignment of feedback profiles and motor readouts
   1.6 Tone input, task timing, and reward
   1.7 Action/No-Action decision
   1.8 TD reward-prediction error
   1.9 Opponent Context→D/P2 plasticity
   1.10 TC plasticity used only in the cue-triggered action task
   1.11 Ablation, plasticity-off, and identity-shuffle controls
   1.12 Simulations and analyses

2. Operant conditioning of neuronal activity (SNOC)
   2.1 Shared circuit and session initialization
   2.2 Reward-free baseline and threshold calibration
   2.3 SNOC reward rule, RPE, and D/P2 plasticity
   2.4 Target definition and session structure

3. Two-alternative forced-choice task
3.1 Modifications to the shared circuit
3.2 Two random cues and intermediate stimuli
3.3 Response timing and outcomes
3.4 Two Go probabilities and a common decision time
3.5 Action-specific eligibility and D/P2 plasticity
3.6 Endpoint training, plasticity freeze, and psychometric testing
3.7 Psychometric analysis

4. Implementation and reproducibility

5. Virtual tracing
5.1 Study design
5.2 CN and SNr source experiments
5.3 Representative source projection images
5.4 VGLUT2 line profile
5.5 CCF streamline-based L1/L5 split
5.6 Single-neuron tracing
5.7 Representative VM neuron
5.8 Bulk anterograde tracing
5.9 Source-to-thalamus weights
5.10 Matched nuclei and two-stage estimation
5.11 Equal-nucleus control
5.12 Final VGLUT2 comparison
5.13 Software and quality control

This Supplementary Information defines the shared BWSLC (Brain-Wide Synergistic Loop Circuit) implementation used for the cue-triggered action task (Go/No-Go task) and SNOC (single-neuron operant conditioning). The baseline implementation is termed syn-BC in the code packages and corresponds to the Full model (BC model) in the main text. SNOC used the same circuit architecture, action/no-action decision rule, TD-RPE, and D/P2 plasticity without pretraining, with task-specific cue and reward procedures. The forced-choice extension retained the shared recurrent and subcortical pathways but modified action representation, decision sampling, and trial-level eligibility.

**Abbreviations:** BG, basal ganglia; CB, cerebellum; TC, thalamocortical; RNN, recurrent neural network; TD-RPE, temporal-difference reward-prediction error; TCP, thalamocortical plasticity; D, direct pathway; ID, indirect pathway; HD, hyperdirect pathway; LTP, long-term potentiation; LTD, long-term depression; SD, standard deviation.

### 1. Cue-triggered action task and shared BWSLC implementation

This section defines the shared cortical–subcortical circuit. Task-specific SNOC and forced-choice procedures are described in Sections 2 and 3.

#### 1.1 Cortical RNN architecture and activity dynamics

The cortical RNN comprised eight channels (modules), with 25 units per channel, for a total of 200 units. Each channel contained 10 excitatory Context units, five excitatory Action units, five excitatory No-Action units, and five inhibitory units. SNOC retained the same population labels and readouts, whereas in the forced-choice task the Action and No-Action populations were reinterpreted as the Go and No-Go populations for each action. Neural states were updated at 1-ms intervals.

Let $v_i(t)$ denote the internal state and $r_i(t)$ the nonnegative firing rate. The total input was

$$u_i(t) = b_i + \sum_j W_{ij}\, r_j(t) + I_i^{\text{cue}}(t) + I_i^{\text{rew}}(t) + I_i^{\text{BC}}(t) + \xi_i(t),$$

The state and rate were updated as

$$v_i(t + \Delta t) = v_i(t) + \frac{\Delta t}{\tau_r}\left[-v_i(t) + u_i(t)\right],$$

$$r_i(t) = \text{clip}\left(g_i(t) v_i(t), 0, r_{\text{max}}\right)$$

where $\Delta t = 1$ ms, $\tau_r = 20$ ms, and $r_{\text{max}} = 100$. $g_i(t)$ was the multiplicative gain applied to the motor populations through BG/CB coincidence and was set to 1 for Context and inhibitory units. Each unit received the following sustained background input:

$$b_i = \max(0.02,\ 0.35 + 0.05\varepsilon_i), \qquad \varepsilon_i \sim \mathcal{N}(0,1)$$

The initial state was $v_i(0) = \max(0, b_i + 0.10\zeta_i)$, where $\text{clip}(z, 0, r_{\text{max}}) = \min[r_{\text{max}}, \max(z, 0)]$. Input-current noise was generated as an Ornstein–Uhlenbeck (OU) process with a time constant of 5 ms and a stationary standard deviation of 0.24.

$$\xi_i(t + \Delta t) = a_\xi \xi_i(t) + 0.24\sqrt{1 - a_\xi^2}\,\eta_i(t), \qquad a_\xi = e^{-\Delta t/5\text{ ms}},$$

where $\eta_i(t) \sim \mathcal{N}(0,1)$.

### 1.2 Recurrent connectivity

The recurrent matrix was oriented as $W_{ij} = W_{\text{target } i, \text{source } j}$. The sign of all outgoing connections was fixed according to the presynaptic unit type, imposing Dale's law: connections from excitatory sources were nonnegative and those from inhibitory sources were nonpositive. Self-connections were not allowed. Initial nonzero weight magnitudes were drawn from a half-normal distribution with an added offset of 0.001, before row balancing and spectral-radius normalization. Within-channel connection probabilities were 0.40 for Context–Context, 0.35 for motor–motor, 0.25 for Context–motor, 0.50 for excitatory-to-inhibitory, 0.80 for inhibitory-to-excitatory, and 0.80 for inhibitory-to-inhibitory connections. Across channels, the connection probability was 0.07 for excitatory sources and 0 for inhibitory sources. Every postsynaptic unit was constrained to receive at least one excitatory and one inhibitory input. For each postsynaptic row, the ratio of total inhibitory to total excitatory input was drawn from a log-normal distribution with mean 1.0 and log standard deviation 0.10 and clipped to 0.70–1.30. Finally, the spectral radius of the recurrent matrix was normalized to 1.0. Recurrent connections were fixed in all experiments.

### 1.3 Context activity and six corticofugal pathways

For each Context unit, phasic activity was obtained by subtracting the 10-s slow baseline $\bar{r}_{ch,j}(t)$ from its firing rate.

$$x_{ch,j}(t) = r_{ch,j}(t) - \bar{r}_{ch,j}(t).$$

Six nonnegative readouts, D, ID, HD, P1, P2, and P3, were constructed from the Context population in each channel. Initial weights were drawn from a zero-mean Gaussian distribution, rectified at zero, and scaled so that the L1 norm was identical across pathways and channels. The raw pathway signal was

$$z_{p,ch}(t) = \kappa \sum_{j=1}^{10} w_{p,ch,j}\, x_{ch,j}(t),$$

where $p \in \{D, ID, HD, P1, P2, P3\}$, and the common pathway scale was $\kappa = 5.0$. Each raw signal was convolved with the pathway-specific causal alpha kernel $h_p$,

$$y_{p,ch}(t) = \left(h_p * z_{p,ch}\right)(t)$$

yielding the filtered signal. Filter timing for each pathway was determined by the base delay and the within-kernel shift. Excluding the thalamocortical return, the pathway-specific onset times were 16, 29, 9, 20, 20, and 60 ms for D, ID, HD, P1, P2, and P3, respectively (Table S1). CB normalization and a scale factor of 1.2 were applied to P1–P3, whereas the original BG scaling was retained for D, ID, and HD.

**Table S1. Temporal kernels of the six pathways (excluding the thalamocortical return delay)**

| Pathway | Alpha time constant | Pathway base delay | Within-kernel shift | Integration target | Thalamus→cortex |
|---|---|---|---|---|---|
| D | 8 ms | 5 ms | 11 ms | BG | 20 ms |
| ID | 6 ms | 5 ms | 24 ms | BG | 20 ms |
| HD | 5 ms | 5 ms | 4 ms | BG | 20 ms |
| P1 | 5 ms | 10 ms | 10 ms | CB | 20 ms |
| P2 | 20 ms | 10 ms | 10 ms | CB | 20 ms |
| P3 | 12 ms | 10 ms | 50 ms | CB | 20 ms |

#### 1.4 BG/CB integration, thalamocortical return, and cortical feedback

Before returning to cortex, the six pathways within each channel were integrated as

$$B_{ch}(t) = D_{ch}(t) - ID_{ch}(t) - HD_{ch}(t),$$

$$C_{ch}(t) = P1_{ch}(t) - P2_{ch}(t) + P3_{ch}(t)$$

where $B$ was the merged BG output and $C$ the merged CB output. Under the standard condition, channel mapping was the identity mapping, such that the subcortical output from channel $ch$ was returned to the same cortical channel.

The merged BG and CB outputs were returned to the cerebral cortex through the thalamus. In addition to pathway-specific filter delays, a common thalamocortical return delay of 20 ms was applied to both merged outputs. In the implementation, the delay was introduced as identical 20-ms leading zero padding in all six pathway kernels; because the same delay applied to every term in the linear integration, this was equivalent to delaying the merged BG and CB outputs by 20 ms.

$$d_{TC} = 20\,ms, \quad t_{cortex,p} = t_{onset,p} + d_{TC}$$

Accordingly, the earliest cortical arrival times of the D, ID, HD, P1, P2, and P3 components were 36, 49, 29, 40, 40, and 80 ms, respectively.

Positive and negative components were defined as

$$B_{\mathrm{pos},ch} = \max(B_{ch}, 0), \qquad B_{\mathrm{neg},ch} = \max(-B_{ch}, 0)$$

$$C_{\mathrm{pos},ch} = \max\left(C_{ch}, 0\right), \qquad C_{\mathrm{neg},ch} = \max\left(-C_{ch}, 0\right)$$

The positive components provided additive current to the first motor population, whereas the negative components provided additive current to the second motor population. Under the Full-model condition, the additive gains for both BG and CB were 6.5, and the summed BG+CB additive current for each sign was capped at 2.5. When the cap was reached, the BG:CB ratio was preserved.

Same-sign BG/CB coincidence was defined as

$$M_{\mathrm{pos},ch} = \left(B_{\mathrm{pos},ch}/s_B\right)\left(C_{\mathrm{pos},ch}/s_C\right)$$

$$M_{\mathrm{neg},ch} = \left(B_{\mathrm{neg},ch}/s_B\right)\left(C_{\mathrm{neg},ch}/s_C\right)$$

with $s_B = s_C = 0.13856$. The multiplicative gain applied to the motor populations was

$$G_{s,ch} = 1 + g_{\max}\frac{M_{s,ch}}{1 + M_{s,ch}}, \qquad s \in \{\mathrm{pos}, \mathrm{neg}\}, \quad g_{\max} = 2.0$$

which constrained the gain to the range 1–3. BG/CB signals influenced behavior through cortical RNN activity.

### 1.5 Alignment of feedback profiles and motor readouts

For each of the two five-unit motor populations in each channel, a nonnegative profile vector was drawn from $|\mathcal{N}(0,1)| + 0.2$ and normalized to an L2 norm of 1. The same vector was used both (i) as the spatial profile of additive BG/CB feedback and (ii) as the cortical motor-readout weights. Thus, the direction of cortical activity amplified by feedback was structurally aligned with the direction measured by the behavioral readout. Motor-readout weights remained fixed throughout. Thalamocortical feedback weights were fixed except in the local-calibration and TCP conditions. For local calibration (Fig. 6), correlations between Action-unit activity and action probability were measured during a separate 10-min spontaneous run of the untrained network with the same seed. The calibrated feedback profile was applied at task step 6000 (10 min). Each Action feedback weight was updated by adding 32 times the mean pre-calibration weight multiplied by its activity–action correlation, clipped to nonnegative values, and rescaled to preserve the total Action-feedback L1 norm.

### 1.6 Tone input, task timing, and reward

In the cue-triggered action task, a single tone instructed an action. Tone input was restricted to Context and inhibitory units. Afferents to Context units in each channel were 1.2|N(0,1)|, and those to inhibitory units were scaled by 0.05. The tone was presented once during 50–100 ms of the trial-onset block. At reward delivery, a weak nonnegative reward afferent generated from an independent random-number stream (standard scale, 0.10) was delivered to Context and inhibitory units during 0–50 ms of the outcome block.

The first trial began after a 3000-ms initial wait, and the inter-trial interval (ITI) after each trial was 1000 ms. The valid action-latency window was 100–1000 ms. Actions within this window received reward code 2, whereas invalid actions or no-action responses at the deadline received reward code 1.

#### 1.7 Action/No-Action decision

At each 1-ms time point, fixed readouts were computed from the Action and No-Action populations across all eight channels, and their respective 10-s slow baselines were subtracted.

$$q_{\text{action}}(t) = b_{\text{action}} + \frac{g_m}{\sqrt{8}} \sum_{ch=1}^{8} \left(p_{ch}(t) - \bar{p}_{ch}(t)\right),$$

$$q_{\text{no}}(t) = b_{\text{no}} + \frac{g_m}{\sqrt{8}} \sum_{ch=1}^{8} \left(n_{ch}(t) - \bar{n}_{ch}(t)\right),$$

where $g_m = 0.8$, $b_{\text{action}} = 0$, and $b_{\text{no}} = 1.8$. Within each 100-ms block, a common time point

$$t_{dec} = \underset{t}{\text{argmax}}[q_{\text{action}}(t) - q_{\text{no}}(t)]$$

was selected, and $q_{\text{action}}(t_{dec})$ and $q_{\text{no}}(t_{dec})$ were read from the same RNN state. The action probability was

$$P(\text{action}) = \max\{0.003, \sigma(1.5[q_{\text{action}}(t_{dec}) - q_{\text{no}}(t_{dec}) - 0.5])\}$$

An action or no-action response was then selected by Bernoulli sampling. For the TD-RPE in the cue-triggered action task, $c(\text{action}) = 0.5$ and $c(\text{no-action}) = 0$; $V_t$ was the value $q_{\text{action}}$ or $q_{\text{no}}$ corresponding to the sampled action.

#### 1.8 TD reward-prediction error

Behavioral state was updated in 100-ms steps. The common temporal-difference reward-prediction error (TD-RPE) was

$$\delta_t = (R_{t+1} - 1)r_s - c(a_t) + \gamma V_{t+1} - V_t,$$

$$\gamma = 0.5^{100/3000} = 0.977,$$

where $R$ was the reward code (0, 1, or 2), $r_s = 10$ was the reward scale, $V_t$ was the task-specific cortical action value, and $c(a_t)$ was the action cost. Thus, reward codes 2, 1, and 0 corresponded to TD reward terms of $+10$, 0, and $-10$, respectively. Task-specific action costs and value definitions are described below.

#### 1.9 Opponent Context→D/P2 plasticity

Plasticity was restricted to nonnegative weights from Context to D and P2. ID, HD, P1, P3, cortical recurrent weights, and motor readouts were fixed. Context phasic activity was averaged within each 100-ms block and centered by subtracting a 10-s eligibility baseline. In the cue-triggered action task and SNOC, reward-driven updates used the centered eligibility from the immediately preceding 100-ms block (lag 1).

$$E\_ch, j(t) \ = \ e\_ch, j(t \ - \ 1 \ behavior \ block)$$

The TD-RPE was multiplied by the eligibility vector normalized by its L2 norm, and the resulting updates were accumulated over each 1-s weight-update interval.

$$H\_ch, j \ \leftarrow \ H\_ch, j \ + \ \eta \ \delta\_t \ E\_ch, j \ / \ sqrt(\Sigma\_ch, j \ E\_ch, j^2), \quad \eta \ = \ 0.003$$

Every 1 s, D and P2 weights were updated in complementary directions. Independent Gaussian weight noise ε (SD, 0.001) was added at the same update time.

$$w\_D, ch, j \leftarrow max[0, w\_D, ch, j + 0.15\,(H\_ch, j + \varepsilon\_D, ch, j)]$$

$$w\_P2, ch, j \leftarrow max[0, w\_P2, ch, j - H\_ch, j + \varepsilon\_P2, ch, j]$$

After each update, D and P2 were renormalized separately so that the L1 norm in each channel returned to its initial value, and the hold was reset to zero. When H>0, D underwent LTP and P2 underwent LTD, shifting both B=D−ID−HD and C=P1−P2+P3 in the positive direction. When H<0, D underwent LTD and P2 underwent LTP, shifting the merged feedback in the negative direction. Thus, bidirectional LTP and LTD were implemented as opponent D/P2 updates.

### 1.10 TC plasticity used only in the cue-triggered action task

In the Full model, the thalamocortical feedback profiles returning to cortex were fixed, and plasticity was restricted to D/P2. BC+TCP retained all components of the Full model and additionally made the cell-wise gains $\theta^{BG}_{ch,j}$ and $\theta^{CB}_{ch,j}$ from positive additive BG/CB feedback to Action units plastic. Both gains were initialized to 1.0 and constrained to 0–2.5.

TCP eligibility was defined as the product of the positive additive BG or CB current in the preceding block, the positive phasic activity of the Action unit, the fixed feedback/readout profile, and local gain and postsynaptic gates. The postsynaptic activity gate served as a proxy for dendritic-spike-dependent plasticity. For $X \in \{BG, CB\}$,

$$r_{\mathrm{phasic},ch,j} = \max(\tilde{r}_{ch,j}, 0)$$

$$e^{X}_{ch,j} = I^{X}_{\mathrm{pos},ch} f^{\mathrm{action}}_{ch,j} r_{\mathrm{phasic},ch,j} G_{\mathrm{gain}}\left(G_{\mathrm{pos},ch}\right) G_{\mathrm{post}}\left(r_{\mathrm{phasic},ch,j}\right)$$

the gate was

$$G(z; \vartheta, s) = \min\left(1, \max\left(0, \frac{z - \vartheta}{s}\right)\right)$$

For the gain gate, $\vartheta = 1.15$ and $s = 0.20$; for the postsynaptic gate, $\vartheta = 0.015$ and $s = 0.03$. Eligibility was L2-normalized across BG and CB, all channels, and all Action units.

LTP was applied on action trials with positive RPE, whereas LTD was applied to invalid actions with negative RPE.

$$N_e = \sqrt{\sum_{X,ch,j} \left(e^{X}_{ch,j}\right)^2}$$

$$\Delta\theta^{X}_{ch,j} = 0.001125\,\delta_{\mathrm{pos}} e^{X}_{ch,j}/N_e - 0.00825\,\delta_{\mathrm{invalid}} e^{X}_{ch,j}/N_e$$

Here, $\delta_{\mathrm{pos}} = \max(\delta, 0)$, whereas $\delta_{\mathrm{invalid}} = \max(-\delta, 0)$ was nonzero only for invalid actions. Updates were applied every 1 s, as for D/P2. A weak positive-feedback-to-no-action branch was initialized to 0.02, but its learning rate was set to zero in the selected BC+TCP condition. TCP L2 decay was zero.

This TC plasticity was introduced only in the BC+TCP condition of the cue-triggered action task. All TC learning rates were set to zero and feedback profiles were fixed in SNOC and the forced-choice task.

### 1.11 Ablation, plasticity-off, and identity-shuffle controls

The same set of controls was applied to both the Full model and BC+TCP. For ablations, specified circuit components were set to zero; for plasticity-off controls, the corresponding weights were fixed. In identity-shuffle controls, the amplitude and temporal waveform of the BG/CB signals were preserved, whereas the identity of the Action/No-Action population and cell-wise profile on which the specified feedback stream acted in cortex was replaced with a seed-fixed random profile. The standard channel correspondence was retained.

**Table S2. Control conditions used in the cue-triggered action task**

| Category | Condition | Operation |
|---|---|---|
| Baseline | Full model | Additive feedback, multiplicative gain, and D/P2 plasticity were all enabled. TC plasticity was also enabled in BC+TCP. |
| Feedback form | Linear only | Multiplicative gain was disabled (gain_max_increment = 0); additive feedback was retained. |
| Feedback form | Gain only | Additive BG and CB gains were set to zero; only coincidence gain was retained. |
| Feedback form | No feedback | Both additive feedback and multiplicative gain were disabled. |
| Plasticity off | D plasticity off | Context→D weights were fixed; P2 plasticity was retained. |
| Plasticity off | P2 plasticity off | Context→P2 weights were fixed; D plasticity was retained. |
| System ablation | BG output off | D, ID, and HD were masked before merging, eliminating BG output B. |
| System ablation | CB output off | P1, P2, and P3 were masked before merging, eliminating CB output C. |
| Identity shuffle | All identity shuffle | The identities of both additive and gain feedback were randomized. |
| Identity shuffle | BG identity shuffle | The identity of BG-derived feedback was randomized. |
| Identity shuffle | CB identity shuffle | The identity of CB-derived feedback was randomized. |
| Identity shuffle | Additive identity shuffle | Only the additive feedback profile was randomized. |
| Identity shuffle | Gain identity shuffle | Only the identity of the population receiving multiplicative gain was randomized. |

#### 1.12 Simulations and analyses

Each condition of the Full model and BC+TCP was run in 100 independently seeded networks (seeds 91–190). Each run comprised 12,000 behavioral blocks (1200 s, 20 min) and was parallelized across 20 workers. All controls were evaluated using the same seed set and task parameters.

Learning curves were aggregated in 10-s bins, and the numbers of rewarded and invalid actions were converted to rates per minute. Endpoint performance was calculated from the final 2 min of each seed and summarized as the across-seed mean and standard error of the mean (SEM).

### 2. Operant conditioning of neuronal activity (SNOC)

#### 2.1 Shared circuit and session initialization

SNOC used the Full model defined for the cue-triggered action task. The cortical RNN, six pathways, BG/CB integration, 20-ms thalamocortical return delay, additive and gain-modulating feedback, action/no-action decision rule, TD-RPE, and opponent Context→D/P2 plasticity were identical to those in Section 1.

Each seed defined an independently initialized network, whose initial state was shared by five target-set sessions. The first 2 min served as a reward-free baseline without tones or externally defined task rewards, immediately followed by 20 min of SNOC. Tone input was zero throughout both baseline and SNOC, and the standard tone–action state machine and task reward were disabled. The same weak reward afferent described in Section 1 was delivered to cortex only when an SNOC reward occurred.

#### 2.2 Reward-free baseline and threshold calibration

D/P2 plasticity was disabled during baseline. For each 100-ms bin, the maximum 1-ms activity of target cell i within the bin, m_i(t), was calculated, and the condition-specific SNOC decision signal was defined as follows.

$$m_i(t) = \max_{\tau \in bin(t)} r_i(\tau)$$

$$s_{single}(t) = m_{target}(t)$$

$$s_{two}(t) = m_{target\,1}(t) - m_{target\,2}(t)$$

$$s_{six}(t) = \sum_{i=1}^{3} m_i(t) - \sum_{i=4}^{6} m_i(t)$$

Signal values observed during baseline were used as candidate thresholds. For each candidate threshold θ, an event was counted when s(t)>θ and at least 1 s had elapsed since the previous virtual event; the number of events N(θ) during the 2-min baseline was then calculated.

$$\theta_{opt} = \arg\min_{\theta \in S_{baseline}} |N(\theta) - 20|$$

Thus, the session-specific threshold θopt that yielded the event count closest to 10 events/min (20 events in 2 min) during baseline was selected. If the signal remained above threshold, another event could occur after the 1-s refractory interval.

#### 2.3 SNOC reward rule, RPE, and D/P2 plasticity

The 20-min SNOC period began immediately after baseline. At each 100-ms step, one reward was delivered when the following condition was satisfied.

$$R_{SNOC,t} = \mathbb{1}\left[s(t) > \theta_{opt} \ \wedge \ t - t_{last\ reward} \geq 1\ s\right]$$

SNOC reward depended only on the target neural signal. The cortical RNN nevertheless continued to use the action/no-action decision rule from Section 1, and an action cost of 0.5 was applied to spontaneous actions in the TD-RPE. Reward code 2 was assigned at SNOC reward delivery and reward code 1 otherwise.

D/P2 plasticity was re-enabled at SNOC onset. The RPE, lag-1 centered Context eligibility, 1-s hold application, opponent D/P2 update, nonnegativity constraint, and within-channel L1 renormalization were identical to those in Sections 1.8–1.9. Updates were driven by the eligibility of all Context inputs and the global TD-RPE.

#### 2.4 Target definition and session structure

Targets were sampled without replacement from each seed's action populations across the five sessions, yielding disjoint cells, pairs, or sets. The 100 sessions comprised 20 independent network seeds with five target sets per seed; sessions sharing a seed were not treated as independent networks in the seed-level analyses.

Table S3. SNOC conditions and session structure

| Condition | Signal | Target composition | Seeds | Sessions per seed | Total sessions |
|---|---|---|---|---|---|
| single target | s_single | One action-related cell | 121–140 | 5 cells per seed | 100 |
| two-target difference | s_two | target 1 − target 2 | 101–120 | 5 disjoint pairs per seed | 100 |
| six-target difference | s_six | Sum of targets 1–3 − sum of targets 4–6 | 141–160 | 5 disjoint sets per seed | 100 |

### 3. Two-alternative forced-choice task

#### 3.1 Modifications to the shared circuit

The forced-choice experiments retained the shared Full-model RNN, six pathways, BG/CB integration, additive and gain-modulating feedback, and opponent D/P2 plasticity, while TCP was disabled. Channels 0–3 were assigned to action 1 and channels 4–7 to action 2. The original Action population in each channel served as the Go population for the action assigned to that channel, and the original No-Action population served as the corresponding No-Go population. Both-Go and neither-Go outcomes were treated as no-action.

#### 3.2 Two random cues and intermediate stimuli

Cue 1 and Cue 2 were independent signed Gaussian vectors $\mathbf{u}_1$ and $\mathbf{u}_2$ restricted to Context and inhibitory units in all eight channels. Each component targeting a Context or inhibitory unit was drawn independently from

$$u_{k,i} \sim \mathcal{N}(0, 5.0^2), \qquad k \in \{1,2\},$$

For the signed interpolation coordinate $c \in [-1,1]$, the input vector was

$$\mathbf{I}(c) = \frac{1-c}{2}\mathbf{u}_1 + \frac{1+c}{2}\mathbf{u}_2$$

Here, $c = -1$ corresponded to pure Cue 1, $c = +1$ to pure Cue 2, and $c = 0$ to an equal-weight mixture. The cue began 50 ms after trial onset and the same vector was presented continuously until an action was selected or the trial timed out.

**3.3 Response timing and outcomes**

The first decision was made at 200 ms after trial onset. Following a no-action outcome, the cue remained on and the decision was repeated every 100 ms. Response opportunities occurred at 200, 300, …, and 1100 ms; the trial timed out if an exclusive Go response was never obtained. The initial wait was 3000 ms and the ITI was 1000 ms.

During training, endpoint Cue 1 (correct action, action 1) and Cue 2 (correct action, action 2) were presented with equal probability. Correct choices, incorrect choices, and timeouts received reward codes 2, 1, and 0, respectively, corresponding to TD reward terms of $+10$, 0, and $-10$. In test trials with $c = 0$, the sensory input was always the equal-weight mixture, whereas the rewarded action alone was randomized 50:50 across trials.

**3.4 Two Go probabilities and a common decision time**

At each 1-ms time point, the 10-s slow baselines were subtracted from the fixed Go and No-Go readouts in channel $ch$, yielding

$$d_{ch}(t) = g_m\left[\left(p_{ch}(t) - \bar{p}_{ch}(t)\right) - \left(n_{ch}(t) - \bar{n}_{ch}(t)\right)\right], \qquad g_m = 0.8,$$

The evidence for each four-channel action group was square-root-normalized in the same manner as the original population readout.

$$e_1(t) = \frac{1}{\sqrt{4}}\sum_{ch=0}^{3} d_{ch}(t), \qquad e_2(t) = \frac{1}{\sqrt{4}}\sum_{ch=4}^{7} d_{ch}(t).$$

Within each 100-ms decision block, the original single-time-point peak detector was extended symmetrically to two actions. A single common time point

$$t_{dec} = \operatorname*{argmax}_t \max\left(e_1(t), e_2(t)\right)$$

was selected, and the following values were read from the same RNN state:

$$q_1 = e_1(t_{dec}), \qquad q_2 = e_2(t_{dec})$$

The Go probability for each action was calculated using the same logistic function.

$$p_a = \sigma[1.5(q_a - 0)], \qquad a \in \{1,2\}.$$

Independent Bernoulli samples $g_a \sim \mathrm{Bernoulli}(p_a)$ were generated. Action 1 was selected when $(g_1, g_2) = (1,0)$, and action 2 when $(0,1)$. The outcomes $(1,1)$ and $(0,0)$ were treated as no-action, and the trial continued.

3.5 Action-specific eligibility and D/P2 plasticity

At each decision attempt $k$, the Bernoulli policy scores were

$$s_{1,k} = g_{1,k} - p_{1,k}, \qquad s_{2,k} = g_{2,k} - p_{2,k}$$

The score-function factor follows the REINFORCE/policy-gradient approach used in reward-based RNN training (Song et al., 2017). The TD value signal was defined as in Section 1.8.

Let $e_{ch,j}^{(k)}$ denote the centered Context eligibility in the 100-ms block. Trial-level eligibility was accumulated as

$$E_{ch,j} \leftarrow E_{ch,j} + s_{a(ch),k} e_{ch,j}^{(k)},$$

across all decision attempts, where $a(ch) = 1$ for $ch = 0, \ldots, 3$ and $a(ch) = 2$ for $ch = 4, \ldots, 7$; the trace decay was 1.0. Both-Go and neither-Go were both behaviorally classified as no-action, but they contributed to eligibility because they had different policy scores.

The TD-RPE obtained at the terminal outcome was multiplied by trial eligibility, and the same D/P2 update as in Section 1.9 was applied. In the forced-choice task, a common cost of 0.5 was applied symmetrically to actions 1 and 2, and the cost of no-action was zero. For an exclusive choice, $V_t$ was the $q_a$ of the selected action; for no-action, it was zero.

#### 3.6 Endpoint training, plasticity freeze, and psychometric testing

Each simulation comprised 20,000 behavioral blocks (2000 s, 33.3 min). During the first 12,000 blocks (1200 s, 20 min), D/P2 was trained using only endpoint cues with $c = \pm 1$. At block 12,000, D/P2 updates, pathway-weight noise, and L1 renormalization were completely stopped. During the remaining 8,000 blocks (800 s, 13.3 min),

$$c \in \{-1, -0.75, -0.5, -0.25, -0.125, 0, 0.125, 0.25, 0.5, 0.75, 1\}$$

were presented in random order. The primary analysis used 100 networks with independent seeds 1–100, run in parallel across 20 workers.

3.7 Psychometric analysis

At each coherence, using only trials with an exclusive choice as the denominator, we calculated

$$P(A_2 \mid c, \text{choice}) = \frac{N(A_2, c)}{N(A_1, c) + N(A_2, c)}$$

Timeout rate, mean decision time, number of decision attempts, number of both-Go conflicts, and number of neither-Go outcomes were quantified separately.

Choice data were fitted with a logistic function containing lower and upper lapse parameters,

$$P(A_2 \mid c) = \lambda_L + (1 - \lambda_L - \lambda_U) \frac{1}{1 + \exp[-k(c - c_0)]}$$

using Bernoulli maximum likelihood. $c_0$ was the midpoint of the logistic core (equal to the point of subjective equality when the two lapse rates are equal), and $k$ was the slope. The lapse parameters were constrained only by

$$\lambda_L \geq 0, \qquad \lambda_U \geq 0, \qquad \lambda_L + \lambda_U < 1$$

Optimization used L-BFGS-B from multiple initial values. The primary curve was fitted after pooling choice trials across all 100 seeds; seed-specific curves and the PSE distribution were treated as secondary analyses.

### 4. Implementation and reproducibility

The model was implemented in Python using NumPy, SciPy, Numba, pandas, PyYAML, and Matplotlib. Neural states and weights were computed primarily in float32, whereas eligibility, TD-RPE, and selected accumulators were computed in float64. Neural updates used a 1-ms time step; behavioral state, decisions, RPE, and eligibility were updated primarily in 100-ms

steps. Random seeds ensured reproducibility of network initialization, dynamic noise, and independent afferent streams.
The cue-triggered action simulations using the Full model and BC+TCP corresponded to source package bc_tcp_synbc_summary_20260710, the SNOC campaign to snoc_no_pretraining_operant_package_20260711, and the psychometric model to bc_grouped_go_forced_choice_v6_20260721. Each package contained the configuration, execution scripts, and analysis scripts used. The consolidated repository separates these engine snapshots under models/bc, models/snoc, and models/forced_choice, and provides portable BSVT/BBVT reanalysis under tracing. Compact checkpoints support numerical reproduction without rerunning the full campaigns.

**Table S4. Major shared parameters and task-specific settings**

| Parameter | Cue-triggered action | SNOC | Forced choice |
|---|---|---|---|
| Neural step / rate time constant | 1 ms / 20 ms | Same as action task | Same as action task |
| Channels × units | 8 × 25 | Same as action task | Same as action task |
| Context / motor 1 / motor 2 / I | 10 / Action 5 / No-Action 5 / 5 | Same as action task | 10 / Go 5 / No-Go 5 / 5 |
| Recurrent spectral radius | 1.0 | 1.0 | 1.0 |
| OU noise SD / time constant | 0.24 / 5 ms | Same as action task | Same as action task |
| Cue input | Nonnegative tone; Context scale 1.2, inhibitory scale 0.05 | tone = 0 | Two independent signed Gaussian vectors; SD 5.0, inhibitory scale 1.0 |
| Cue timing | Once at 50–100 ms after trial onset | None | Continuous from 50 ms until action or timeout |
| Decision onset | Valid actions from 100 ms onward | Original decision rule at 100-ms intervals | First choice at 200 ms |
| ITI / initial wait | 1000 / 3000 ms | No trial task; SNOC reward refractory period, 1000 ms | 1000 / 3000 ms |
| β / motor gain | 1.5 / 0.8 | Same as action task | 1.5 / 0.8 |
| Additive BG/CB gains / cap | 6.5 / 6.5 / 2.5 | Same as action task | Same as action task |

| Parameter | Cue-triggered action | SNOC | Forced choice |
|---|---|---|---|
| Coincidence scales / max increment | 0.13856 / 0.13856 / 2.0 | Same as action task | Same as action task |
| TD half-life / reward scale | 3000 ms / 10 | Same as action task | Same as action task |
| Action cost | 0.5 for action only | 0.5 for spontaneous action only | 0.5 for both actions 1 and 2 |
| D/P2 learning rate / D scale | 0.003 / 0.15 | Same as action task | Same as action task |
| Pathway update / weight-noise SD | 1000 ms / 0.001 | Same as action task | Same as action task (frozen during testing) |
| TC plasticity | Enabled only in the BC+TCP condition | OFF | OFF |
| Duration | 20 min | 2 min baseline + 20 min SNOC | 20 min training + 13.3 min test |

### 5. Virtual tracing

#### 5.1 Study design

We combined source-to-thalamus input distributions from the cerebellar nuclear group not classified as inhibitory-specific (CN) or the substantia nigra pars reticulata (SNr) with nucleus-specific thalamocortical output distributions to estimate relative cortical-layer profiles for the CN→thalamus→cortex and SNr→thalamus→cortex pathways. Thalamocortical outputs were quantified independently using Brain Image Library (BIL) single-neuron reconstructions from the BRAIN Initiative Cell Census Network (BICCN) and bulk anterograde tracing data from the Allen Mouse Brain Connectivity Atlas. Thalamic-nucleus abbreviations are listed in Extended Data Table 1.

#### 5.2 CN and SNr source experiments

CN comprised six experiments in which the dentate nucleus (DN) was a primary or secondary injection structure and the experiment was not inhibitory-specific. The SNr group comprised seven retained experiments. Let $p_{e,n}$ denote the structure-unionize projection density for experiment e and thalamic nucleus n, and $J_e^{\mathrm{src}}$ the source injection volume. The mean source-group input density was: Source unionizes used hemisphere_id=3 (bilateral) and is_injection=False; experiment IDs, transgenic lines, and primary/secondary injection structures are listed in data/tracing/source_experiments.csv.

$$u_{s,n} = \frac{1}{N_s}\sum_{e\in s} \frac{p_{e,n}}{J_e^{\mathrm{src}}}$$

For the pattern-only heatmap, each source column was divided by its maximum:

$$h_{s,n} = \frac{u_{s,n}}{max_{n'} u_{s,n'}}$$

Column-scaled values were used only to display nucleus patterns and not to compare absolute source magnitudes.

#### 5.3 Representative source projection images

The representative CN case was DN-primary Slc17a6-IRES-Cre experiment 286608092, and the representative SNr case was SNr-primary Gad2-IRES-Cre experiment 303534443. The Allen 50-μm registered projection-density volume $P_e(v)$ was divided by injection volume:

$$g_e(v) = \frac{P_e(v)}{J_e}$$

Signal was summed within the Common Coordinate Framework (CCF) thalamus mask $T_k$ for each coronal plane k, and the plane with the maximum signal was selected:

$$k_e^* = argmax_k \sum_{v\in T_k} g_e(v)$$

Display contrast was normalized independently by the 99.5th percentile $Q_{e,0.995}$ of nonzero thalamic values and square-root transformed:

$$R_e(v) = \sqrt{clip(\frac{g_e(v)}{Q_{e,0.995}}, 0,1)}$$

This transformation was used only for the representative images.

#### 5.4 VGLUT2 line profile

A linear fluorescence profile $I(x)$ was acquired from the pia toward the white matter in an M1 sagittal section stained for vesicular glutamate transporter 2 (VGLUT2). The original display used manually specified bins: L1a 10–25, L1b 26–50, L2/3 51–105, L5a 106–130, L5b 131–200, L6a 201–270, and L6b 271–340. Mean intensity in bin $B_l$ was:

$$\bar{I}_l = \frac{1}{|B_l|} \sum_{x \in B_l} I(x)$$

These original bins were treated separately from the final streamline-depth split described below.

**5.5 CCF streamline-based L1/L5 split**

We used the Allen CCF 100-µm annotation (Wang et al., 2020a) and ccf-streamlines. Each cortical voxel v was assigned to its nearest streamline. Relative depth was calculated from the pial-side start of the parent layer $a_{k,0}$, the white-matter-side end $a_{k,1}$, and the streamline position $s_v$:

$$r_v = \frac{s_v - a_{k,0}}{a_{k,1} - a_{k,0}}$$

For L1 and L5, $r_v$<0.5 was classified as upper and $r_v$≥0.5 as lower. Allen annotations for L2/3, L4, L6a, and L6b were retained. Voxels without a valid streamline lookup retained the previous distance-based assignment as a fallback. The same mask was applied to the single-neuron and bulk datasets. Relative depths from -0.05 to 1.05 were accepted and clipped to [0,1] before splitting. These operational upper/lower halves were not assumed to be identical to cytoarchitectonic sublayers; fallback counts are provided in mask_qc.csv.

**5.6 Single-neuron tracing**

We analyzed neuronal-morphology files in SWC format from 195 BIL/BICCN thalamic neurons. SWC coordinates were expressed in CCF micrometers and used a scale factor of 1.0; type 2 was treated as axon, and segments were resampled at 10-µm intervals. Let $L_{i,a,l}$ be the axon length for cell i, area a, and layer l, and $V_{a,l}$ the area–layer volume. Mean density for soma nucleus n was: Segment midpoint samples were assigned to 100-µm voxels by rounding CCF coordinates divided by 100 to the nearest integer (NumPy rint); bilateral area-layer volumes were used.

$$d_{n,a,l}^{\text{single}} = \frac{1}{N_n} \sum_{i \in n} \frac{L_{i,a,l}/1000}{V_{a,l}}$$

Cell–area–layer combinations with no detected axon were explicitly zero-filled before averaging.

**5.7 Representative VM neuron**

For each VM neuron i, we calculated the number of M1/M2/S1 target areas containing nonzero L1 axon, $A_i$, and the total L1 axon length across target areas, $L_i^{\text{L1}}$:

$$A_i = |\{a : L_{i,a,\text{L1}} > 0\}|$$

$$L_i^{\text{L1}} = \sum_a (L_{i,a,\text{L1upper}} + L_{i,a,\text{L1lower}})$$

Lexicographic ranking first by the area count and then by $L_i^{\text{L1}}$ was used to select cell 1119751254_18458_5057-X16469-Y15624_reg. Type-2 segment midpoints were assigned to the split mask and displayed in sagittal projection.

**5.8 Bulk anterograde tracing**

We analyzed 103 experiments with primary thalamic injection sites. For experiment e, projection-density volume $P_e(v)$ was averaged within area–layer mask $M_{a,l}$ and divided by injection volume $J_e$: Projection density denotes the fraction of detected projection pixels, not mean fluorescence intensity. The archived calculation did not apply the Allen data_mask; valid-voxel-only reaggregation is provided as a sensitivity option.

$$d_{e,a,l}^{\text{bulk}} = \frac{1}{J_e}\frac{1}{|M_{a,l}|}\sum_{v\in M_{a,l}} P_e(v)$$

Values were then averaged across experiments with the same primary nucleus n, yielding $d_{n,a,l}^{\text{bulk}}$. Injection-volume normalization was performed before averaging across experiments.

**5.9 Source-to-thalamus weights**

Given estimated neuron number $C_n$, the source input weight was:

$$w_{s,n} = C_n u_{s,n}$$

This quantity is a relative weight proportional to the estimated neuron number, not an absolute synapse count. Cn was calculated as nucleus volume in the 25-μm CCF annotation multiplied by the corresponding Blue Brain Cell Atlas neuron-density estimate. The numerical densities, region matches, fallback labels, and resulting counts are archived in neuron_estimates.csv.

5.10 Matched nuclei and two-stage estimation

The primary comparison was restricted to 12 nuclei represented in both methods (Allen ontology acronyms): CM, IMD, PIL, PO, PVT, RE, RT, SMT, VAL, VM, VPL, and VPMpc. For method m: Nuclei without single-neuron output profiles, including PF, PCN, and SPFp, were excluded rather than assigned zero output. The final streamline-based analysis used all 195 available cells, without the earlier 15-cell-per-nucleus subsampling.

$$D_{s,a,l}^{(m)} = \sum_{n\in T} w_{s,n} d_{n,a,l}^{(m)}$$

M1 was MOp, M2 was MOs, and S1 comprised SSp-bfd, SSp-ll, SSp-m, SSp-n, SSp-tr, SSp-ul, and SSp-un. Multi-subarea groups A were volume-weighted:

$$D_{s,A,l}^{(m)} = \frac{\sum_{a\in A} D_{s,a,l}^{(m)} V_{a,l}}{\sum_{a\in A} V_{a,l}}$$

Layer fractions were normalized within each method, source, and area:

$$F_{s,A,l}^{(m)} = \frac{D_{s,A,l}^{(m)}}{\sum_{l'\in\mathcal{L}_A} D_{s,A,l'}^{(m)}}$$

**5.11 Equal-nucleus control**

For the VGLUT2 comparison (Extended Data Fig. 2d), the equal-nucleus control used the arithmetic mean of output densities from the 12 matched thalamic nuclei, with a weight of 1/12 for each nucleus: This equal-nucleus control differs from the equal-weight average of the CN and SNr source-group profiles; significance values from source-experiment sign-flip tests were not assigned to the equal-nucleus control.

$$d_{\text{equal},a,l}^{(m)} = \frac{1}{|T|}\sum_{n\in T} d_{n,a,l}^{(m)}$$

Here, the subscript equal denotes equal weighting, and T comprised the 12 nuclei listed in Section 5.10. The same area aggregation and layer normalization were applied; the resulting profiles were used for the VGLUT2 comparison in Section 5.12.

#### 5.12 Final VGLUT2 comparison

Background b was the mean over profile indices 0–6, and $I'(x) = max(0, I(x) - b)$. For the final comparison, manually defined parent-layer ranges were divided into nonoverlapping depth halves, and background-subtracted layer means were normalized within area: Inclusive final bins were M1: 10–29, 30–50, 51–105, 106–152, 153–200, 201–270, and 271–340; M2: 7–27, 28–48, 49–100, 101–140, 141–180, 181–280, and 281–340 (L1 upper, L1 lower, L2/3, L5 upper, L5 lower, L6a, L6b). L4 was absent. These are profile indices, not calibrated physical depths.

$$G_{A,l} = \frac{\bar{I}'_{A,l}}{\sum_{l' \in \mathcal{L}_A} \bar{I}'_{A,l'}}$$

Agreement with VGLUT2 was summarized by the mean absolute error (MAE), calculated as the mean absolute difference across the seven valid layers: MAE was expressed in percentage points by multiplying this fraction-scale value by 100. Each measured line profile was treated as a fixed reference, without estimation of between-animal staining variability.

$$\mathrm{MAE}_A = \frac{1}{|\mathcal{L}_A|} \sum_{l \in \mathcal{L}_A} |F_{A,l} - G_{A,l}|$$

#### 5.13 Software and quality control

Analyses used Python 3.10, NumPy, pandas, SciPy, Matplotlib, AllenSDK, h5py, pynrrd, and ccf-streamlines. We recorded the number of out-of-bounds SWC samples, streamline coverage, fraction-sum errors, experiment counts, and cell counts. Lines, bars, text, and axes were stored as vector objects in PDF files; only heatmap matrices and representative projection-density images were rasterized.
